# Dermatopontin-expressing fibroblasts mediate an essential skin macrophage niche

**DOI:** 10.1101/2024.11.21.624708

**Authors:** Apple Cortez Vollmers, Sunny Z. Wu, Anthony Altieri, Erika E. McCartney, Hannah Bender, Wyne P. Lee, Juan Zhang, Crystal Hu, Salil Uttarwar, Jason A. Vander Heiden, Christopher Davidson, Yein Chung, Willie Ortiz, Michael Long, Raymond Asuncion, Yeqing Angela Yang, Jean X. Jiang, Zora Modrusan, Akshay T. Krishnamurty, Wenxian Fu, Sören Müller, Matthew B. Buechler, Shannon J. Turley

## Abstract

Fibroblasts are present in all tissues and are crucial for maintaining tissue homeostasis. We previously identified fibroblasts marked by *Dermatopontin* (*Dpt*) but their role in supporting macrophage homeostasis remains unclear. Here, we generated novel mesenchymal lineage-restricted genetic tools to target *Dpt* expressing fibroblasts and elucidate their role in supporting skin macrophages. Transcriptional profiling, flow cytometry, and in situ hybridization uncovered two broad populations of F4/80-expressing skin macrophages, denoted by high expression of CD206 and CD64 (CD206^hi^CD64+), or CD11c. Targeted depletion of *Dpt*+ fibroblasts resulted in a profound loss of both macrophage populations. Conditional deletion of colony-stimulating factor-1 (*Csf1)* in *Dpt*+ fibroblasts revealed that CD206^hi^CD64+, and not CD11c+, macrophages are acutely dependent on fibroblast-derived *Csf1*, consistent with their higher expression of the *Csf1* receptor. Following *Csf1* deletion in *Dpt*+ fibroblasts, loss of CD206^hi^CD64+ macrophages were observed across the dermis, dermal white adipose tissue (dWAT), and adventitia, accompanied by a modest upregulation of fibroblast-related and extracellular matrix (ECM) genes and structural changes to the skin. Alterations to the skin network upon loss of fibroblast-derived *Csf1* and CD206^hi^CD64+ macrophages led to a significant delay in wound healing. We also demonstrate the CSF1-CSF1R signaling pathway is functionally relevant in human systemic sclerosis, or scleroderma, as elevated levels of CSF1 produced by fibroblasts and an increased abundance of macrophages both correlate with disease severity. Our findings demonstrate the role of *Dpt*+ fibroblasts in regulating a *Csf1*-dependent macrophage niche in skin and orchestrating responses in injury and disease.

## Introduction

The skin serves as a protective barrier and is composed of various cell types, including fibroblasts and macrophages. Fibroblasts define tissue architecture by producing and remodeling ECM proteins^1^, and provide a supportive niche for different cell populations^2–4^. Macrophages are key effector cells of the immune system. Both cell types are integral in maintaining tissue homeostasis and coordinating response to injury and disease^2^.

Macrophages reside in different layers of the skin. While fibroblast-macrophage interactions have been explored in healthy^5^ and injured skin^6–8^, the function of specific fibroblast populations in supporting macrophages across compartments of healthy skin is largely unknown. Moreover, fibroblasts may mediate the action of macrophages in skin wound healing and repair^6–9^ but the precise transcriptional types of fibroblasts and macrophages that orchestrate these processes remains unclear.

Previous studies have demonstrated that homeostatic maintenance of macrophages in many tissues depends on colony-stimulating factor-1 (CSF1)^5,10–13^, a growth factor crucial for macrophage proliferation, differentiation and maintenance. Fibroblasts have been proposed to provide a key source of CSF1 to maintain macrophage homeostasis^5,10–13^. In the skin, the cellular source of CSF1, and contribution of fibroblast-derived factors to support macrophage development and homeostasis are incompletely understood.

Here, we sought to investigate a potential cell circuit between macrophages and fibroblasts in the skin. *Dermatopontin (Dpt)*-expressing fibroblasts have been suggested to operate as resource cells, giving rise to specialized fibroblasts across different tissues during development, steady-state, and in disease^14^. While the abundance of *Dpt*-expressing cells varies across tissues, in the skin, a high proportion of fibroblasts (>80%) show robust expression of *Dpt*^14^. The high representation of *Dpt* expression among fibroblasts raised the possibility that *Dpt+* fibroblasts exhibit functions in the skin aside from strictly operating as a resource cell for differentiating fibroblasts, and explored the hypothesis that homeostasis of skin macrophages is regulated by *Dpt*+ fibroblasts.

By implementing novel genetic mouse models combined with single-cell transcriptomics analysis, we demonstrate that depletion of *Dpt+* fibroblasts leads to a marked reduction in two major populations of skin macrophages marked by CD206^hi^CD64+ or CD11c+. Cell signaling predictions between *Dpt*+ fibroblasts and both macrophage populations revealed *Csf1*-*Csf1r* signaling as one of the top five most significant interactions. To test this prediction, we confirmed that the majority of fibroblast subtypes in skin express *Dpt* and are the main source of CSF1. Next, we generated a new mouse model to conditionally knock out *Csf1* in *Dpt+* fibroblasts and found CD206^hi^CD64+ macrophages to be solely dependent on fibroblast-derived CSF1 in skin. The resulting loss of CD206^hi^CD64+ macrophages, which are found in dermis, dWAT and adventitia, were necessary to facilitate an effective wound healing response. Importantly, we demonstrate that the CSF1-CSF1R axis is relevant in human disease, as patients with scleroderma, an autoimmune disease with inflammatory and fibrotic processes^15^, exhibit increased levels of fibroblast-derived *CSF1* and macrophage abundance which correlate with disease severity.

## Results

### Targeted depletion of *Dpt*+ fibroblasts affects multiple skin fibroblast transcriptional states

To examine the individual subtypes of cells in skin, single-cell sequencing using cellular indexing of transcriptomes and epitopes (CITE-Seq)^16^ was performed on fluorescence activated cell sorting (FACS)-sorted CD45-EPCAM-CD31-PDPN+ fibroblasts and CD45+ immune cells from the skin of *Dpt^IresCreERT^*^2^*;Rosa26*^lox-stop-loxYFP^ mice (i*Dpt;R26*^YFP-CTRL^)^14^. These mice were given the estrogen analog tamoxifen to elicit *Cre* recombination such that *Dpt*-expressing cells acquired yellow fluorescent protein (YFP). Initial analysis of the fibroblast compartment identified five broad transcriptional populations in steady state skin marked by 1) *Pi16+*, 2) *Pi16*+*Col15a1*+, 3) *Cthrc1*+, 4) *Cxcl12*+, and 5) *Sparc*+. (Fig. 1a, top). As insertion of the *CreERT2* construct at the 3’ end of *Dpt* prevents direct detection by scRNA-Seq, the YFP reporter in i*Dpt;R26*^YFP/DTR^ mice was utilized as proxy for *Dpt* expression using a custom genome alignment. *Yfp* was found to be expressed across all fibroblast populations (Fig. 1a, bottom, Extended data Fig.1a).

**Figure 1.**
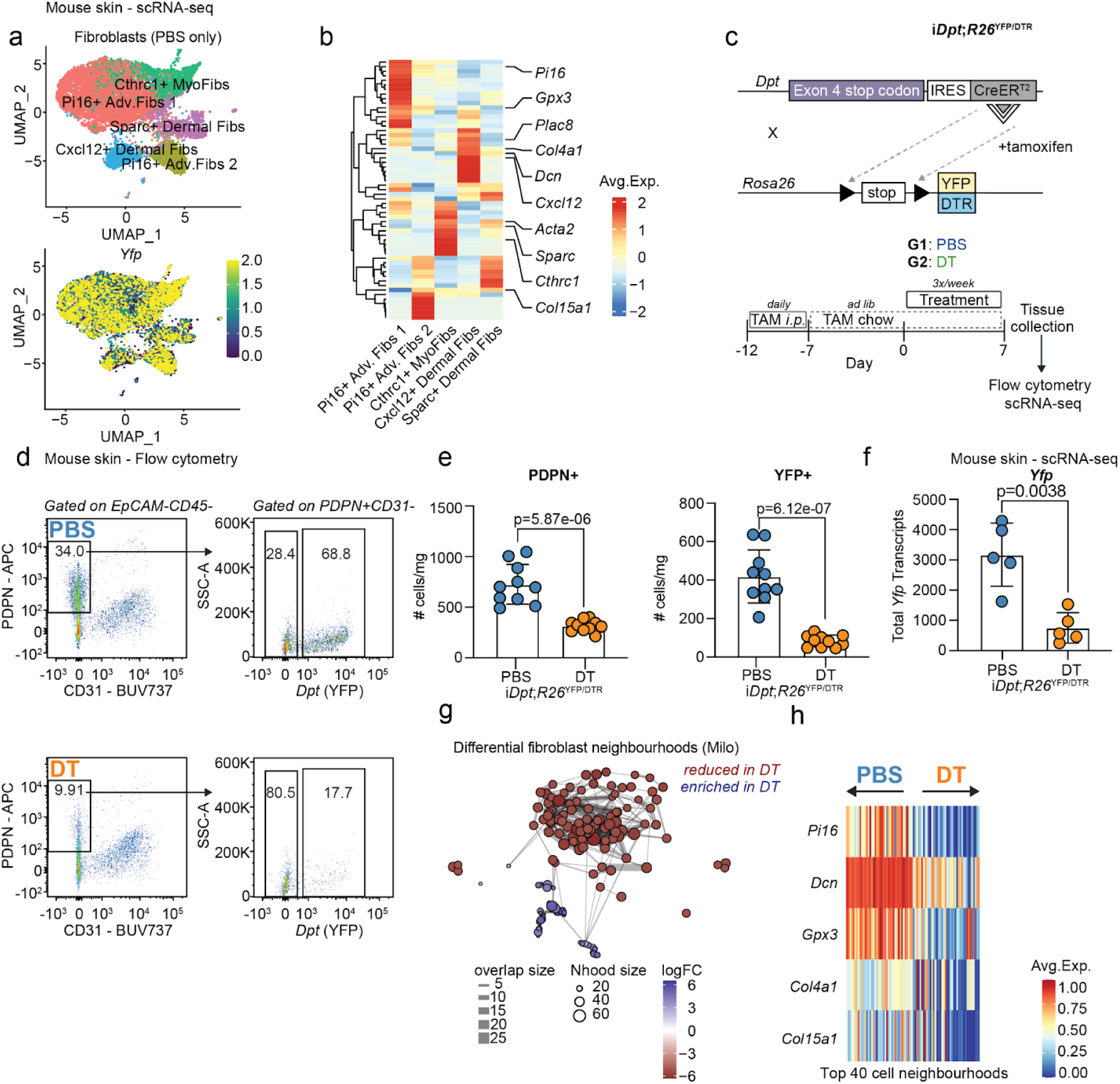
Targeted depletion of *Dpt+* fibroblasts. **a,** Single-cell sequencing in steady-state skin tissue of i*Dpt;R26*^YFP-CTL^ mice (*n*=5, representative of 1 independent experiment). Five skin fibroblast populations (top) identified based on canonical marker genes. Feature plot (bottom) of expression values for *Yfp*. **b,** Cluster averaged heatmap log-normalized gene expression of fibroblast-related genes used for cluster annotation. **c,** Schematic of the genetic (top) and experimental approach (bottom) for the generation of i*Dpt;R26*^YFP/DTR^ mice. Mice were administered with tamoxifen for approximately two weeks, and then treated 3X/week with either PBS or diphtheria toxin (DT). **d,** Flow cytometry analysis of live, CD45-EpCAM-PDPN+CD31-cells in the skin of i*Dpt;R26*^YFP/DTR^ mice treated with PBS (*n*=10) and DT (*n*=10). Representative flow cytometry plot showing the frequency of PDPN+ and YFP+ fibroblasts. Representative of 7 independent experiments. **e,** Quantification of total number of PDPN+CD31-cells (left) and total number of YFP+ cells (right). Bar plots represent Mean ± SD. Statistics were calculated using unpaired, two-tailed, Student’s *t*-test. **f,** Expression of unique *Yfp* transcripts from scRNA-seq analysis performed on fibroblast cells showing statistically significant enrichment in PBS samples compared to DT. Summed counts for *Yfp* transcripts are shown. Statistics were calculated using unpaired, two-tailed, Student’s *t*-test. **g,** Differential abundance of fibroblast neighbourhoods across PBS or DT conditions. Analysis performed using the Milo package in R. Significant neighbourhoods (-logFDR<0.05) enriched in PBS or DT conditions are indicated by red or blue, respectively. **h,** Expression of selected fibroblast marker genes across differentially abundant fibroblast neighbourhoods. Top 40 cell neighbourhoods enriched in each group are shown (-logFDR<0.05). Positive and negative fold changes reflect cell neighbourhoods enriched in DT (arrow pointing right) vs PBS (arrow pointing left) groups, respectively.

Building on previously published work, these five fibroblast populations were further characterized by identifying their markers, including those indicating spatial location within skin^17^. The two clusters of *Pi16*+ fibroblasts showed differential abundance of *Col15a1* and co-expression of *Gpx3*, a marker of fibroblasts that spatially locate to the hypodermal/adventitial region of the skin^17^*. Cthrc1*+ fibroblasts expressed *Acta2* (α-SMA), a marker of myofibroblasts. *Cxcl12*+ and *Sparc*+ fibroblasts expressed *Dcn*, a marker that spatially locates to the dermis^17^, with differential expression of *Plac8*, *Col4a1*, and *Col15a1* (Fig. 1b).

To investigate the role of fibroblasts in supporting cutaneous macrophages in vivo, a genetic mouse model was generated to deplete *Dpt*+ fibroblasts, which represent the majority of fibroblasts in skin (Fig. 1a-c)^2,14,18^. i*Dpt;R26*^YFP-CTRL^ animals were bred to animals harboring a *diphtheria toxin* receptor (DTR) cassette on the other R26 allele^14^ to generate i*Dpt;R26*^YFP/DTR^ mice (Fig. 1c). i*Dpt;R26*^YFP/DTR^ mice were treated with tamoxifen to activate *Cre* and induce YFP and DTR expression followed by PBS (control) or *diphtheria toxin* (DT) administration to deplete *Dpt*+ fibroblasts. Seven days after the start of PBS or DT treatment, full-thickness samples of skin were collected from the flank for analysis (Fig. 1c).

Flow cytometry was performed to quantify total loss of YFP-expressing cells. YFP expression was restricted to the CD45-EpCAM-CD31-PDPN+ (PDPN+) fibroblast compartment (Extended data Fig. 1b) and marked approximately 60% of PDPN+ fibroblasts (Fig. 1d), consistent with previous reports^14^. Fibroblasts, as measured by enumerating non-hematopoietic, non-endothelial PDPN+ (Fig. 1e, left) or YFP+ cells (Fig. 1e, right), were significantly reduced upon DT administration, confirming the validity of this model.

A significant decrease in *Yfp* transcripts was shown by scRNA-seq following DT treatment, consistent with flow cytometry profiling and confirming successful depletion of *Dpt*+ fibroblasts at the transcriptional level (Fig. 1f). Striking differences between PBS and DT treated animals (Fig. 1g, h) were revealed by analysis of the fibroblast compartment. To identify cell neighborhoods with differential abundance following depletion of *Dpt*+ fibroblasts, Milo analysis was applied to single-cell data^19^ (Fig. 1g, Extended data Fig. 1c,d). Decreased expression of *Pi16*, *Dcn, Col15a1*, *Gpx3* and *Col4a1,* markers of fibroblast clusters across different spatial compartments within skin^17^ (Fig. 1b), was shown in fibroblast neighborhoods most significantly reduced after depletion of *Dpt*+ fibroblasts (indicated to the right; Fig. 1h). These data suggest *Dpt*+ fibroblasts encompass multiple populations in dermis and adventitia, and their depletion may in turn compromise neighboring cells dependent on them for their maintenance.

### *Dpt*+ fibroblasts support the homeostasis of two major skin macrophage subsets

To identify the broad transcriptional changes in skin tissue following DT-induced *Dpt+* fibroblast depletion, differential expression analysis between pseudo-bulk profiles of all cell types from PBS and DT-treated samples was performed. An enrichment of fibroblast-related genes and pathways in the PBS condition, including *Pi16*, *Col1a1* and *Fn1*, and pathways related to ECM-related gene ontologies was revealed (Fig. 2a). Furthermore, an enrichment of immunoregulatory pathways associated with myeloid cell migration, activation and homeostasis was identified (Fig. 2a), suggesting depletion of *Dpt*+ fibroblasts may result in a loss of a myeloid-supporting niche. *Csf1* is a central feature of these myeloid-related pathways (Fig. 2a), and in DT-treated samples, a reduction in normalized pseudo-bulk expression of *Csf1* was observed (Fig. 2b). Expression of *Csf1* across different cell types in skin was evaluated and fibroblasts were identified as the primary cellular source of this growth factor (Fig. 2c), consistent with previous findings^5^. Since fibroblasts in skin do not express *Csf1r* (Fig. 2c), these data suggest that fibroblasts provide pro-survival and trophic factors, such as *Csf1,* in a paracrine manner to support skin-resident macrophages.

**Figure 2.**
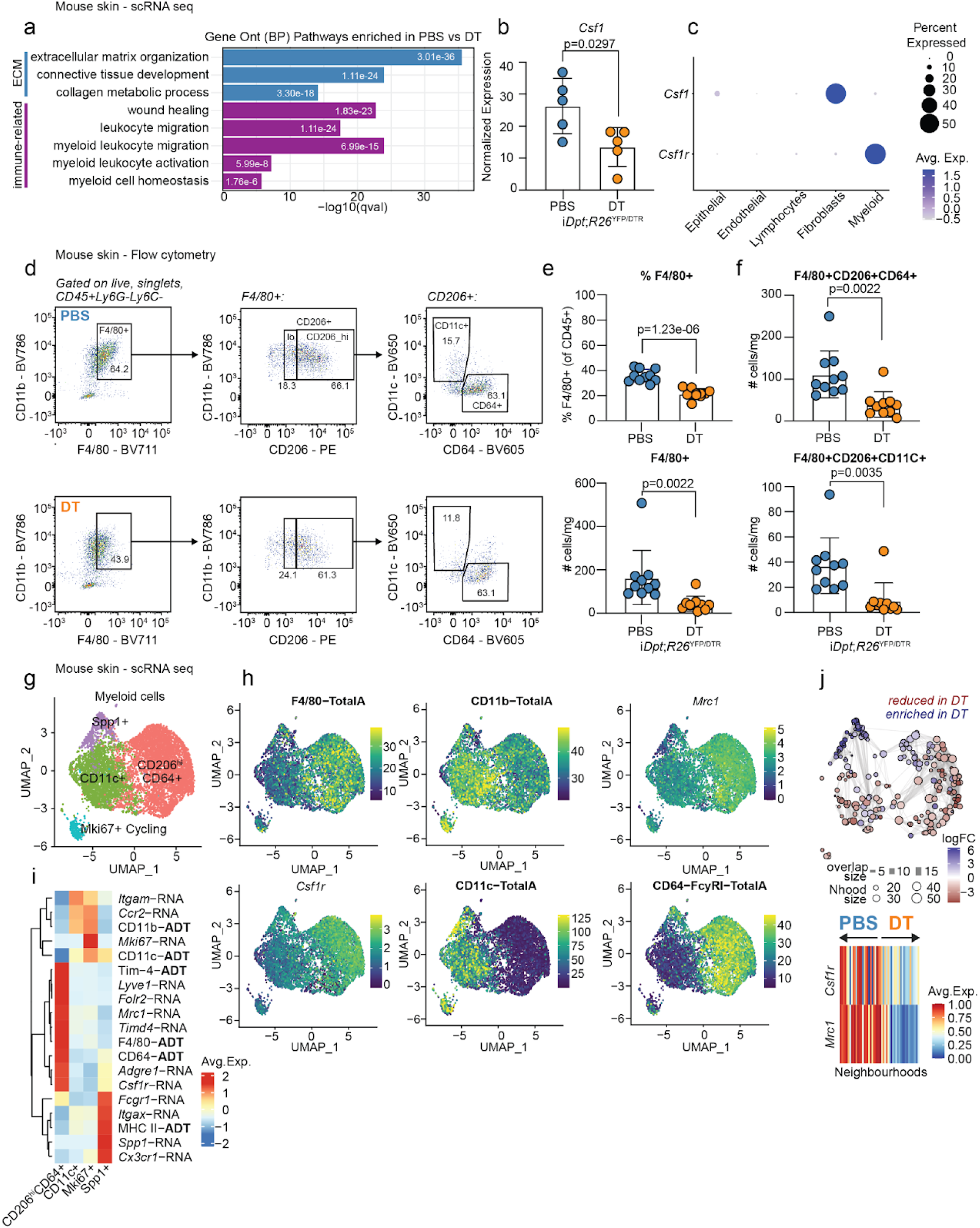
*Dpt+* fibroblast depletion results in CSF1 and macrophage reduction in skin. **a,** scRNA-seq analysis of mouse skin tissue (flank) from i*Dpt;R26*^YFP/DTR^ mice. Selected biological pathways enriched in PBS (*n*=5) compared to DT-treated (*n*=5) skin tissue samples, as identified using pseudo bulk differential expression analysis. All significant genes identified using DESeq 2 were used for enrichment in the Gene Ontology Biological Processes database. **b,** Expression of *Csf1* showing statistically significant enrichment in PBS samples compared to DT. Summed counts for unique Normalized pseudo bulk expression values are shown for *Csf1.* **c,** Expression of *Csf1* and *Csf1r* across different cell types sampled in skin tissue. Cell types annotated using SingleR with the ImmGen reference. **d,** Flow cytometry analysis of live, CD45+Ly6G-Ly6C-CD11b+ cells in the skin of i*Dpt;R26*^YFP/DTR^ mice treated with PBS (*n*=10) and DT (*n*=10). Representative flow cytometry plot showing the frequency of F4/80+ cells and CD206+, CD11c+ and CD64+ subsets. Representative of 7 independent experiments. **e,** Quantification of percent CD11b+F4/80+ cells (top) and total number of CD11b+F4/80+ cells (bottom). Bar plots represent Mean ± SD. Statistics were calculated using unpaired, two-tailed, Student’s *t*-test. **f,** Quantification of total number of F4/80+CD206+CD64+ (top) F4/80+CD206+CD11c+ (bottom) cells. **g,** UMAP visualization of myeloid cells from scRNA-seq performed on skin from i*Dpt;R26*^YFP/DTR^ mice (*n*=5 control, *n*=5 DT-treated, representative of 1 independent experiment). **h,** Feature plot of expression values for F4/80, CD11b, *Mrc1*, *Csf1r*, CD11c, and CD64 markers. **i,** Cluster averaged heatmap log-normalized gene expression and antibody-derived tags of macrophage-related markers used for cluster annotation. **j,** Differential abundance of macrophage neighborhoods across PBS or DT conditions (top). Analysis performed using the Milo package in R. Significant neighborhoods (-logFDR<0.05) enriched in PBS or DT conditions are indicated by red or blue, respectively. Decreases in the average expression of *Csf1r* and *Mrc1* (CD206) across Milo cell neighborhoods enriched in DT compared to PBS conditions (bottom). The top and bottom 30 neighborhoods are shown.

To determine the impact of *Dpt+* fibroblast depletion on macrophages, flow cytometry was performed to measure changes amongst F4/80+ macrophages in skin (Fig. 2d). In *Dpt+* fibroblast-depleted mice, both the frequency and total number per milligram of tissue of CD11b+F4/80+ cells were significantly reduced (Fig. 2e). To better distinguish subsets of myeloid cells affected by *Dpt+* fibroblast-depletion, additional markers including CD206, CD11c, and CD64 were used^6,20,21^. With this combination of markers, two major populations were identified that both expressed CD206, but differed in expression of CD11c, an integrin, and CD64, a Fc receptor encoded by *Fcgr1* that mediates binding to IgG antibodies^22,23^. Both, F4/80+CD206+CD11c+ (denoted as CD11c+) and F4/80+CD206+CD64+ (referred hereafter as CD206^hi^CD64+) populations were significantly reduced following *Dpt+* fibroblast-depletion (Fig. 2f, Extended data Fig 1e).

To rule out impact of *Dpt*+ fibroblast depletion on Langerhans and dendritic cell (DC) homeostasis, CD45+EpCAM+CD207+ and CD45+Ly6G-Ly6C-F4/80-MHCII^hi^CD11c^hi^ cells in PBS and DT-treated mice were enumerated, and comparable numbers were found between groups (Extended data Fig 1f). Collectively, these data indicate that selective depletion of *Dpt*+ fibroblasts results in a significant reduction in two populations of F4/80-expressing macrophages, which suggests a role for *Dpt*+ fibroblasts in maintaining skin macrophage homeostasis.

To validate and gain deeper insight into the tissue-specific nature of macrophage loss following *Dpt*+ fibroblast deletion, the myeloid compartment from skin of PBS and DT-treated mice was analyzed using CITE-seq. Here, two broad macrophage populations were identified (Fig. 2g) with distinct gene (RNA) and surface marker (antibody derived tag, ADT) expression (Fig. 2h,i), consistent with our flow cytometry gating strategy. This included an *Mrc1*^hi^ (CD206^hi^) subset that was further defined by F4/80^hi^, CD11b^lo^, *Csf1r*^hi^, *Timd4^hi^*, *Lyve1*+, *Folr2*+, CD11c-, and CD64+, markers representative of tissue resident macrophages^13,24^, and a CD11c+ subset that was further defined by *Mrc1*^lo^, F4/80^low^, CD11b^hi^, *Csf1r^lo^*, *Timd4^lo^, Lyve1-, Folr2-* and CD64-(Fig. 2h,i). Additionally, smaller clusters of *Spp1*+ inflammatory myeloid cells and *Mki67*+ proliferating macrophages were identified (Fig. 2g,i).

To characterize the CD206^hi^CD64+ and CD11c+ macrophage subsets, previously defined markers of skin macrophages, including Major Histocompatibility Complex class II (MHCII), *Ccr2*, *Cx3cr1, Mrc1,* and *Arg1* were used to infer origin, localization, or function^5–7,20,21,24^. From our analysis, MHCII was shown to have broad expression in skin and could not discriminate between the two primary macrophage populations (Extended Data Fig. 2a,b). Instead, the two populations exhibited differential expression of *Ccr2* (Extended Data Fig. 2a,b), suggesting the *Ccr2*^hi^ population of CD11c+ macrophages are monocyte derived, and the *Ccr2*-population of CD206^hi^CD64+ macrophages are tissue resident. Consistent with previous findings, the majority of macrophages from adult mouse skin did not express *Cx3cr1*^5,20,21^ (Extended Data Fig. 2a,b). As *Mrc1 (*CD206) and *Arg1* have been used to delineate macrophage phenotypes in vitro^25^, *Arg1* expression was assessed and little to no expression was found across macrophage populations (Extended Data Fig. 2a,b).

Importantly, F4/80-expressing CD206^hi^CD64+ and CD11c+ macrophage populations did not express *Zbtb46, Xcr1* and *Clec9a* (Extended Data Fig. 2c,d). CD11c+CD64-populations have previously been categorized as dermal DCs^5,20,21^. This population of cells was shown by our analysis to not express markers of DCs (Extended Data Fig. 2c,d).

To further aid in the description of these macrophages, *Mrc1*^hi^ (CD206^h^) macrophages were identified by pathway enrichment analysis as enriched for gene-ontology terms related to regulation of epithelial cells, vasculature development, smooth muscle proliferation, wound healing and phagocytosis, whereas CD11c+ macrophages were enriched in pathways related to cell-cell and cell-matrix adhesion and T-cell activation (Extended Data Fig. 1g).

To further characterize changes to macrophages following *Dpt*+ fibroblast-depletion, Milo differential neighborhood analysis was performed which enabled identification of macrophage cell neighborhoods with differential abundance. Consistent with earlier findings by flow cytometry, abundance of macrophage cell neighborhoods within both CD206^hi^CD64+ and CD11c+ populations was significantly reduced following *Dpt*+ fibroblast-depletion (Fig. 2j, Extended Data Fig. 1h). Consistently, a decrease in *Mrc1* (CD206) expression in macrophage neighborhoods that decreased upon *Dpt+* fibroblast-depletion (Fig. 2j, bottom) was found. In line with the concept that some macrophage states may be regulated by fibroblast-derived CSF1, a decrease in macrophage neighborhoods enriched for *Csf1r* (Fig. 2j, bottom) was observed. In addition, *Spp1*+ inflammatory myeloid cell neighborhoods were increased following *Dpt+* fibroblast-depletion (Fig. 2j). Pathways related to apoptotic cell clearance were enriched in this *Spp1*+ inflammatory cluster (Extended Data Fig. 1g), which may be related to targeted depletion of fibroblasts using DT. These analyses, employing flow cytometry and deep profiling using single-cell and CITE-seq, highlight the critical role of *Dpt*+ fibroblasts in supporting CD206^hi^CD64+ tissue-resident and CD11c+ monocyte-derived macrophages, and underscore their importance in regulating specific immune pathways as well as maintaining tissue homeostasis.

### *Csf1* derived from *Dpt*+ fibroblasts locally supports CD206^hi^CD64+ macrophages in dermis, dWAT and adventitia

To discover candidate factors mediating fibroblast-macrophage interactions, ligand-receptor communication prediction between steady-state control *Dpt*+ skin fibroblasts and both CD206^hi^CD64+ and CD11c+ skin macrophages was performed using CellChat^26^ (Fig. 3a, Extended Data Table 2). This revealed a total of 57 significant interactions, 50 of which were fibroblast ligands directed towards macrophage receptors (Extended Data Table 1). *Csf1*-*Csf1r* was one of the most significant interactions predicted between *Dpt+* fibroblasts and both macrophage populations, with a greater interaction strength toward the CD206^hi^CD64+ subset (Fig. 3a). Other interactions were categorized in Complement, CCL and SEMA3 pathways (C3-C3ar1, Ccl7-Ccr2, Sema3c-Plxnd1, C3-ITGAX_ITGB2; Fig. 3a).

**Figure 3.**
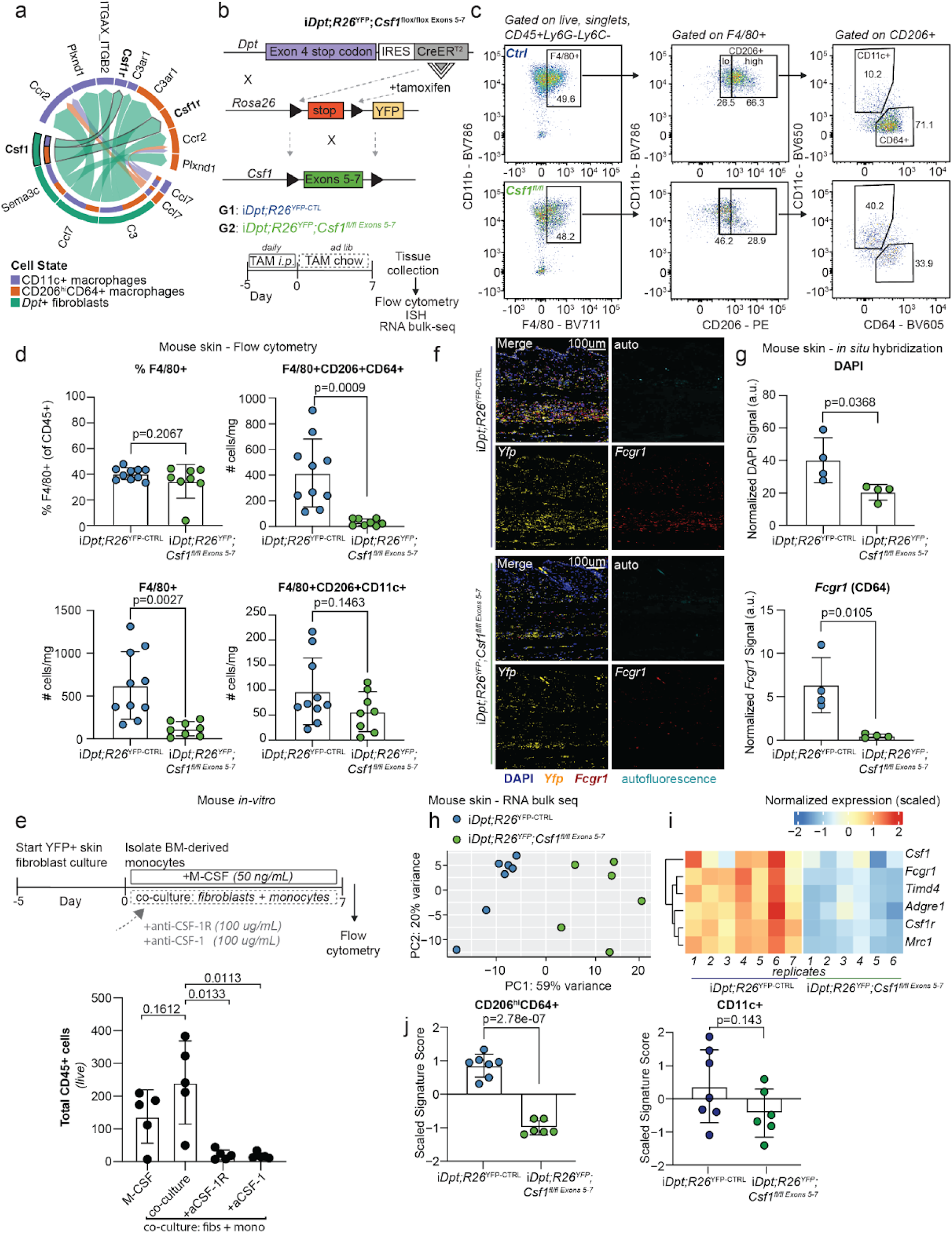
Conditional loss of *Csf1* in *Dpt*+ fibroblasts results in a reduction of CD206^hi^CD64+ skin macrophages. **a,** Circos plot of ligand-receptor interactions between all steady-state fibroblasts (PBS condition) with CD11c+ macrophages or CD206^hi^CD64+ macrophages. **b,** Schematic of i*Dpt;R26*^YFP^;*Csf1^fl/fl Exons^*^5–7^ transgenic mouse. **c,** Representative flow cytometry plot (top) showing the frequency of F4/80+ cells and CD206+, CD11c+ and CD64+ subsets in i*Dpt;R26*^YFP-CTL^ (*n*=10) and i*Dpt;R26*^YFP^;*Csf1^fl/fl Exons^*^5–7^ mice (*n*=8). Representative of 2 independent experiments. Cells were gated on CD45+Ly6G-Ly6C-CD11b+ cells. **d,** Quantification of percent CD11b+F4/80+ cells and total number of CD11b+F4/80+, F4/80+CD206+CD64+ and F4/80+CD206+CD11c+ cells. Bar plots represent Mean ±SD. Statistics were calculated using unpaired, two-tailed, Student’s *t*-test. **e,** Experimental scheme and quantification of flow cytometry data showing total CD45+ cells with addition of CSF-1R and CSF-1 blocking antibodies in each condition. Bar plots represent Mean ± SD. Statistics were calculated using unpaired, two-tailed, Student’s *t*-test. n.d. (not detected). **f,** Representative in situ hybridization (ISH) images for *YFP* (yellow) and *Fcgr1* (red) from the skin of control (*n*=4) and i*Dpt;R26*^YFP^;*Csf1^fl/fl Exons^* ^5–7^ (*n*=4) mice, representative of 1 independent experiment). DAPI signal is shown in blue. Autofluorescence signal is shown in cyan. Scale bars, 100 um. **g,** Quantification of DAPI (top) and *Fcgr1 (*bottom) signal. a.u. stands for arbitrary unit, a unit for intensity. Bar plots represent Mean ± SD. Statistics were calculated using unpaired, two-tailed, Student’s *t*-test. **h,** Principal component analysis (PCA) plot of bulk-RNA sequencing from the skin of i*Dpt;R26*^YFP-CTL^ (*n*=7) and i*Dpt;R26*^YFP^;*Csf1^fl/fl Exons^*^5–7^ (*n*=6) mice. Representative of 1 independent experiment. **i,** Heatmap of log-normalized expression (scaled) for *Csf1* and CD206^hi^CD64+ macrophage marker genes *Adgre1* (F4/80), *Csf1r*, *Fcgr1* (CD64), *Mrc1* (CD206) and *Timd4* across biological replicates of skin tissue isolated from i*Dpt;R26*^YFP-CTL^ (*n*=7) and i*Dpt;R26*^YFP^;*Csf1^fl/fl Exons^* ^5–7^ (*n*=6). **j,** Quantification of single-cell derived gene-signature scores for CD206^hi^CD64+ (left) and CD11c+ (right) macrophages between biological replicates of skin tissue isolated from i*Dpt;R26*^YFP-CTL^ (n=7) and i*Dpt;R26*^YFP^;*Csf1^fl/fl Exons^* ^5–7^ (n=6). The top 50 genes per macrophage cluster were used for mean signature scoring and z-scaling. Statistics were calculated using unpaired, two-tailed, Student’s *t*-test.

To directly test whether provision of CSF1 by *Dpt+* fibroblasts is required to maintain skin macrophages in vivo, *Dpt^IresCreERT2^Rosa26^LSLYFP^Csf1^flox/flox^*mice (i*Dpt;R26*^YFP^;*Csf1*^fl/fl^) were generated that enabled inducible and conditional knockout of *Csf1* in *Dpt*+ fibroblasts upon administration of tamoxifen (Fig. 3b). i*Dpt;R26^YFP^* mice were crossed to two *Csf1* floxed alleles, a novel one with LoxP sites flanking exons 5-7 (i*Dpt;R26*^YFP^;*Csf1^fl/fl Exons^* ^5–7^, Fig. 3b) and a previously published allele with LoxP sites flanking exons 4-6 (i*Dpt;R26*^YFP^;*Csf1^fl/fl Exons^* ^4–6^, Extended data Fig. 4a)^27^ to generate two complementary genetic models to assess the role of *Csf1* provision by *Dpt*+ fibroblasts in skin macrophage homeostasis.

**Figure 4.**
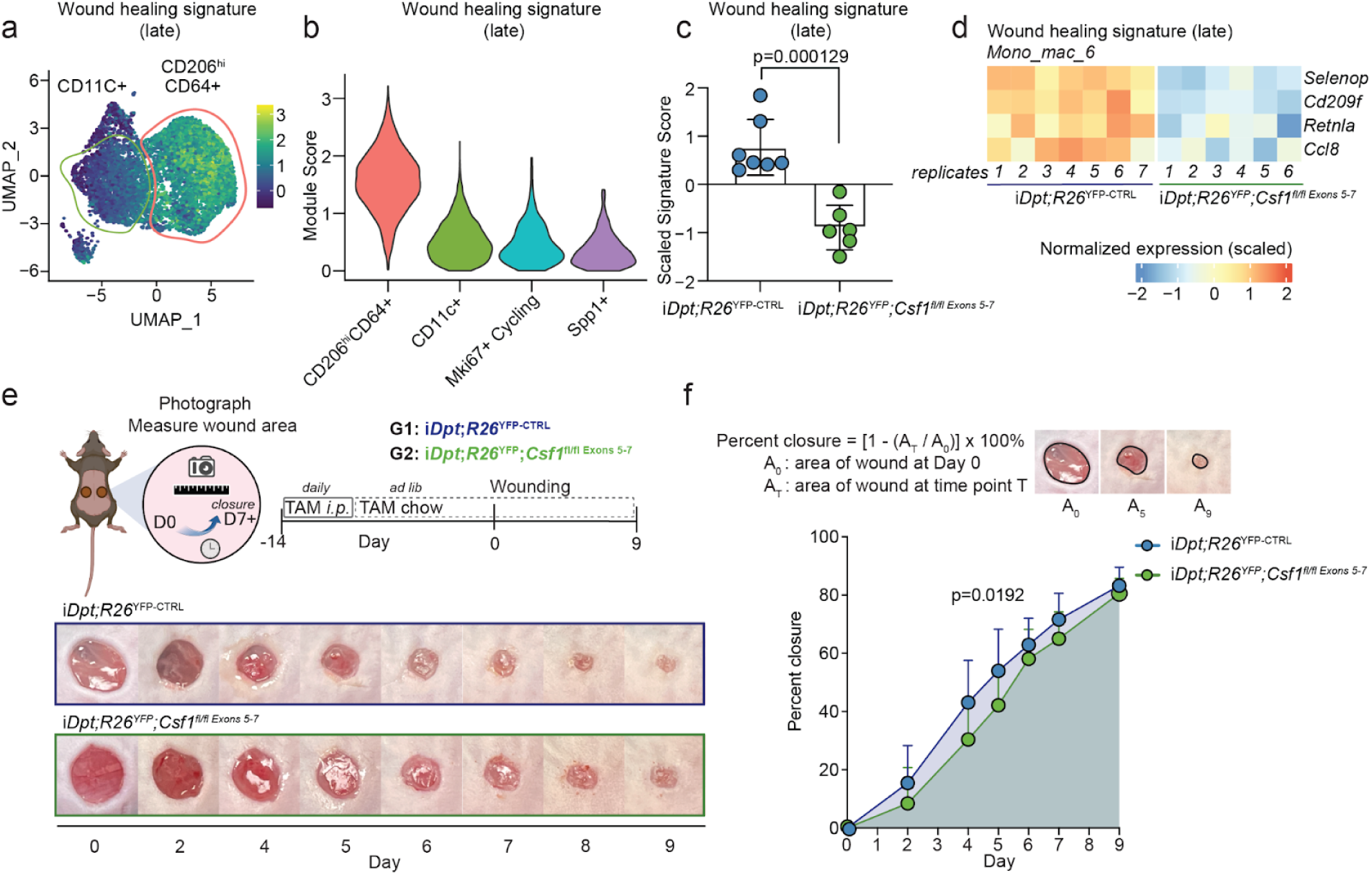
Reduction in macrophages upon loss of fibroblast-derived *Csf1* alters skin wound healing. **a,b,** Featureplot **(a)** and Violin plot **(b)** showing enrichment of a late macrophage wound healing signature^7^ in CD206^hi^CD64+ macrophages. Signature scores were computed using the ModuleScore function in the Seurat package. **c,** Quantification of single-cell derived wound healing gene-signature scores between biological replicates of skin tissue isolated from i*Dpt;R26*^YFP-CTL^ (*n*=7) and i*Dpt;R26*^YFP^;*Csf1^fl/fl Exons^* ^5–7^ (*n*=6) as analyzed by bulk-RNA sequencing. Representative of 1 independent experiment. Statistics were calculated using unpaired, two-tailed, Student’s *t*-test. **d,** Heatmap of log-normalized expression (scaled) for late (characterized as mono-mac-6 population^7^) macrophage wounding healing genes *Selenop*, *Cd209f*, *Retnla* and *Ccl8* across biological replicates of skin tissue isolated from i*Dpt;R26*^YFP-CTL^ (*n*=7) and i*Dpt;R26*^YFP^;*Csf1^fl/fl Exons^* ^5–7^ (*n*=6). Genes selected are statistically significant between groups (p-value<0.05) as determined using DESeq2. **e,** Experimental scheme (top, Created with BioRender) of wound model (*n*=14-19 mice in each group, compiled from two independent experiments). Images of the wound were acquired using a digital camera. Acquired images were analyzed for percent closure on ImageJ/Fiji. Representative images of wounds (bottom) from one i*Dpt;R26*^YFP-CTL^ and i*Dpt;R26*^YFP^;*Csf1^fl/fl Exons^* ^5–7^ mouse are shown. **f,** Quantification of percent closure over time between i*Dpt;R26*^YFP-CTL^ and i*Dpt;R26*^YFP^;*Csf1^fl/fl Exons^* ^5–7^ mice. Line graphs represent Mean ± SD. Statistics were calculated using unpaired, two-tailed, Student’s *t*-test comparing areas under the curve.

Bulk RNA-sequencing was performed on YFP+ fibroblasts isolated from control and i*Dpt;R26*^YFP^;*Csf1^fl/fl Exons^* ^5–7^ mice to confirm the deletion of Exons 5-7 (Extended data Fig. 3a). Next, changes in F4/80-expressing macrophages following conditional knockout of *Csf1* in *Dpt*+ fibroblasts were measured by flow cytometry. A significant decrease in the total number per milligram of tissue of F4/80+ macrophages was observed (Fig. 3c,d, Extended data Fig. 4b-d) in i*Dpt;R26*^YFP^;*Csf1^fl/fl Exons^* ^5–7^ mice. Using markers for CD206, CD11c, and CD64, a significant reduction in the total number of CD206+ cells per milligram of tissue was observed following conditional knockout of *Csf1* in *Dpt*+ fibroblasts (Extended Data Fig. 3b). When changes in CD11c+ and CD206^hi^CD64+ macrophage populations were compared, a marked loss in the total number of CD206^hi^CD64+ cells was observed, with no statistically significant changes in the total number of CD11c+ macrophages per milligram of tissue cells (Fig. 3d). The selective loss of CD206^hi^CD64+ macrophages in i*Dpt;R26*^YFP^;*Csf1*^flox/flox^ mice is aligned with previous scRNA-seq CellChat analysis, which predicted a higher interaction strength between *Dpt*+ fibroblasts and CD206^hi^CD64+ macrophages (Fig. 3a) due to their elevated expression of *Csf1r* (Extended Data Fig. 3c,d). Additionally, the potential for monocyte-derived CD11c+ macrophages to repopulate makes changes in their numbers less noticeable upon *Csf1* conditional knockout in *Dpt+* fibroblasts. Together, our flow cytometry and scRNA-seq data demonstrate that *Csf1r*^hi^ expressing CD206^hi^CD64+ macrophages acutely depend on *Csf1* from *Dpt*+ fibroblasts in vivo.

To support in vivo findings that *Dpt+* fibroblasts and macrophages can directly interact, an in vitro co-culture system was established with *Dpt*+ fibroblasts and macrophages differentiated from bone marrow monocytes. First, CSF1 secretion by *Dpt*+ fibroblasts was confirmed using ELISA (Extended Data Fig. 3e). Next, CSF1 or CSF1R blocking antibodies were added to the co-cultures. A significant reduction in the persistence of F4/80+ macrophages, as measured by total CD45+ cells by flow cytometry, was observed under CSF1 (anti-CSF1) or CSF1R (anti-CSF1R) blocking conditions (Fig. 3e). These results suggest that *Dpt*+ fibroblasts can directly provide CSF1 to support F4/80+ macrophages (Extended Data Fig. 3f-h) via a CSF1/CSF1R signaling dependent mechanism. To further test this hypothesis, fibroblasts isolated from i*Dpt;R26*^YFP^;*Csf1^fl/fl Exons^* ^4–6^ mice were co-cultured directly with GFP-labeled bone-marrow derived macrophages. The loss of *Csf1* in *Dpt*+ fibroblasts led to decreased GFP counts (Extended Data 4e,f), to suggest that fibroblast-derived *Csf1* is critical for macrophage maintenance.

To determine whether the loss of macrophages can be visualized in the skin microenvironment, RNA in situ hybridization (ISH) was performed in mouse skin tissue sections obtained from control and tamoxifen-induced i*Dpt;R26*^YFP^;*Csf1^fl/fl Exons^* ^5–7^ mice (Fig. 3f). *Dpt*+ (*Yfp*) fibroblasts were mapped across multiple layers of skin, including dermis, dWAT, and adventitia, consistent with previous analysis and published data^17^. *Fcgr1* (the gene encoding CD64), was also detected in the dermis, dWAT, and adventitia, indicating fibroblasts and macrophages occupy the same compartments within skin. In i*Dpt;R26*^YFP^;*Csf1^fl/fl Exons^*^5–7^ mice, a marked reduction in *Fcgr1* macrophages in all three compartments was observed, captured both by imaging (Fig. 3f) and quantification of *Fcgr1* signal intensity (Fig. 3g).

As *Dpt*+ fibroblasts are found across tissues, including bone marrow, it was necessary to determine if the reduction of skin macrophages in i*Dpt;R26*^YFP^;*Csf1^fl/fl Exons^* ^4–6^ mice was a consequence of *Csf1* deletion from *Dpt*+ fibroblasts in bone marrow. Bone marrow monocyte frequency was found to be unchanged in i*Dpt;R26*^YFP^;*Csf1^fl/fl Exons^* ^4–6^ mice compared to controls (Extended data Fig. 4g,h). Thus, the reduction in skin macrophages is suggested to result from the absence of local *Dpt*+ fibroblast-derived CSF1 rather than impaired generation in bone marrow. Additionally, to determine if circulating CSF1 regulates macrophage homeostasis in skin, CSF1 levels were measured in serum. No changes in circulating CSF were detected in i*Dpt;R26*^YFP^;*Csf1^fl/fl Exons^* ^4–6^ mice compared to controls (Extended data Fig. 4i). This finding suggests that levels of CSF1 are controlled locally in skin, and *Dpt*+ fibroblast derived CSF1 is required to maintain macrophage homeostasis in skin. In summary, *Dpt*+ fibroblasts in skin are important for maintenance of CD206^hi^CD64+ macrophages found in dermis, dWAT, and adventitia through direct provision of CSF1.

To further profile the transcriptomic differences resulting from the conditional knockout of *Csf1* in *Dpt+* skin fibroblasts, bulk RNA-sequencing was performed on whole-skin isolated from control and i*Dpt;R26*^YFP^;*Csf1^fl/fl Exons^* ^5–7^ mice. Principal Component Analysis (PCA) revealed marked differences in global gene expression profiles between i*Dpt;R26*^YFP-CTRL^ and i*Dpt;R26*^YFP^;*Csf1^fl/fl Exons^* ^5–7^ samples (Fig. 3h), which differential expression analysis revealed to be predominantly driven by macrophage-related genes. A significant reduction was observed in the expression of *Adgre1* (F4/80), *Csf1r, Timd4, Fcgr1* (CD64) and *Mrc1* (CD206) in i*Dpt;R26*^YFP^;*Csf1^fl/fl Exons^* ^5–7^ samples (Fig. 3i). Single-cell derived gene-signature scores for CD206^hi^CD64+ macrophages, but not CD11c+ macrophages (Fig. 3i, Extended data Fig. 3i), were significantly reduced in i*Dpt;R26*^YFP^;*Csf1^fl/fl Exons^*^5–7^ skin, consistent with flow cytometry analysis that found only the frequency and total number of CD206^hi^CD64+ macrophages to be reduced (Fig. 2f). In summary, the i*Dpt;R26*^YFP^;*Csf1*^fl/fl^ mouse models described herein provide in vivo validation that *Dpt*+ skin fibroblasts produce *Csf1* to locally support the maintenance of CD206^hi^CD64+ macrophages.

### Csf1 and macrophage loss affects transcriptional state but not maintenance of skin fibroblasts

A two-cell circuit has previously been modeled in vitro to demonstrate reciprocal exchange of growth factors between fibroblasts and macrophages to control cell numbers^28^. We predicted that loss of *Csf1* in *Dpt*+ fibroblasts would affect not only the homeostasis of macrophages but also impact the fibroblast compartment. To investigate the consequence of CSF1 and macrophage loss on skin fibroblasts in vivo, differences in the abundance of PDPN+ fibroblasts, both by frequency and total number per milligram of tissue, in the skin of i*Dpt;R26*^YFP^;*Csf1^fl/fl Exons^* ^5–7^ mice were measured by flow cytometry. Strikingly, fibroblast abundance was unchanged (Extended data Fig. 5a). By ISH, *Yfp* (*Dpt*) signal intensity was comparable between control and i*Dpt;R26*^YFP^;*Csf1^fl/fl Exons^* ^5–7^ mice (Extended data Fig. 5b). These data suggest that fibroblasts do not rely on growth factors produced by CD206^hi^CD64+ macrophages, and that maintenance of fibroblast numbers in skin are independent of changes in macrophage numbers.

**Figure 5.**
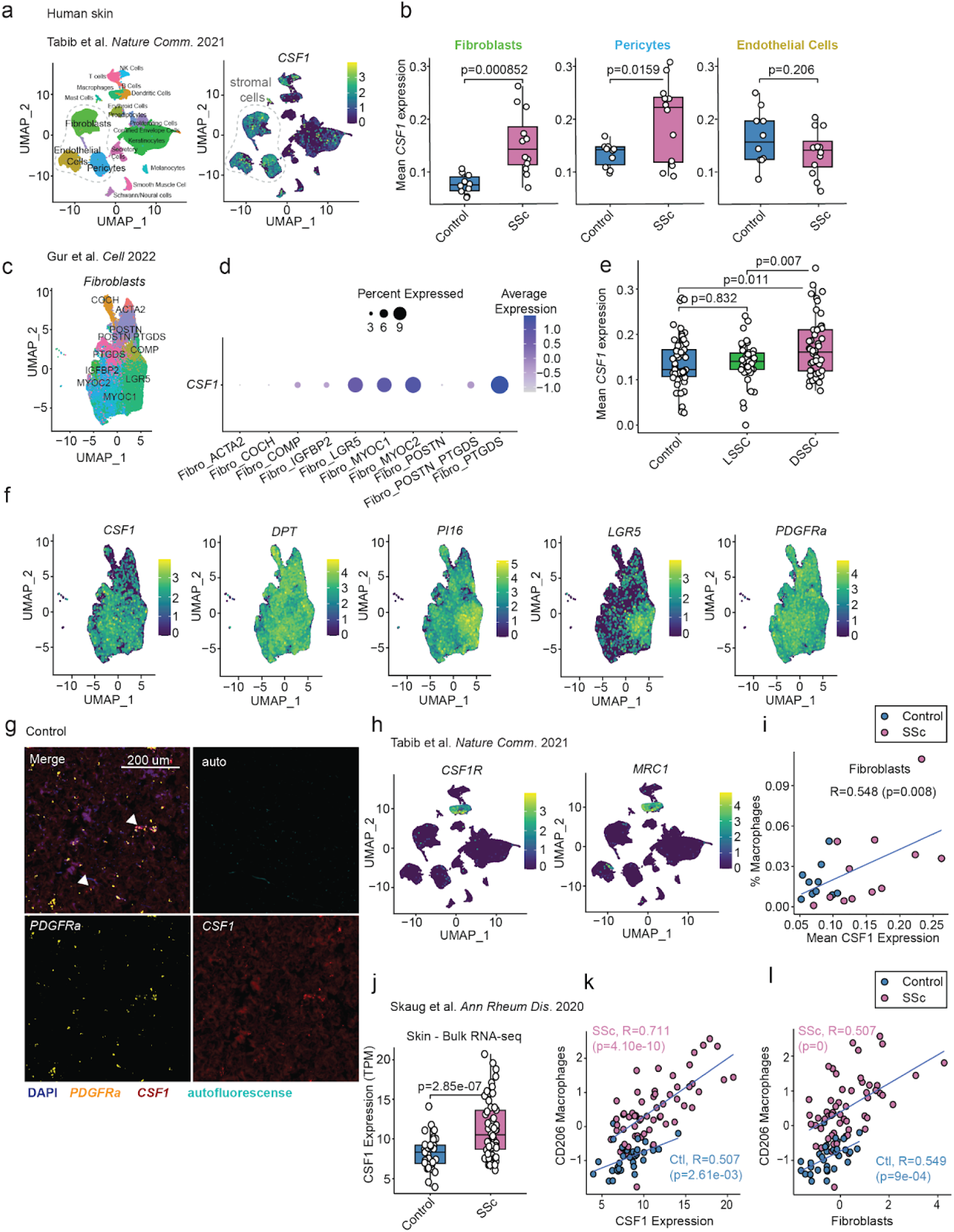
Human skin fibroblast *CSF1* correlates with macrophages. **a,** UMAP (left) representing single-cells from healthy (n=10) and scleroderma, SSc (n=12) human skin tissue . Data re-analyzed from the Tabib 2022 study. Stromal cells including fibroblasts, pericytes, and endothelial cells are indicated. Featureplot (right) showing the expression of *CSF1* across different cell clusters of human scleroderma skin tissue, including enrichment in stromal cells populations. **b,** Quantification of the average *CSF1* expression in fibroblasts, pericytes, and endothelial cells across healthy (n=10) and scleroderma (n=12) patients. For each box-and-whisker plot, the horizontal bar indicates the median. The upper and lower limits of the boxes indicate the first and third quartiles. The ends of the whiskers indicate the min and max. For each box-and-whisker plot, significance was calculated using unpaired, two-tailed Student’s *t*-test. **c,** UMAP representing fibroblasts from healthy control (*n*=56) and SSc patients (*n*=96) from the Gur et al 2022 study^32^. Fibroblast annotations are derived from the original study. **d,** Dot plot visualizing the expression of *CSF1* across human fibroblasts clusters. **e,** Quantification of the average *CSF1* expression in fibroblasts across healthy (blue; *n*=56), limited-SSC (green; *n*=38) and diffuse SSc (pink; *n*=48) patients. **f,** Featureplot showing the expression of *CSF1* and other fibroblast markers, including *DPT, PI16, LGR5, and PDGFRA* across different fibroblast clusters shown. **g,** Representative in situ hybridization (ISH) images for *PDGFRa* (yellow) and *CSF1* (red) from human control skin (n=3) tissue sections from an independent cohort of patients. Representative of 1 independent experiment. DAPI signal is shown in blue. Autofluorescence is shown in cyan. Scale bars, 200 um. **h,** Featureplot showing the expression of *CSF1R* (left) and *MRC1* (right) across different cell clusters of human scleroderma skin tissue, including enrichment in myeloid cell populations. **i,** Correlation plot of the average *CSF1* expression (as in **(e)**) in fibroblasts and the percentage of macrophages of all cells per individual patient. Computed using Pearson correlation. For each Pearson correlation, statistical significance was determined using the *cor.test* function in *R.* Healthy (blue; *n*=10) and scleroderma, SSc (pink; *n*=12) samples are colored. **j,** Normalized expression of *CSF1* (TPM) between SSc (n=55) and healthy control patients (n=33) from the Skaug et al 2019 study^38^. Statistical significance determined using DESeq2. **k,** Pearson correlation between *CSF1* expression and CD206 macrophage gene signature scores in healthy control and SSc samples. **l,** Pearson correlation between CD206 macrophage and fibroblast gene signature scores in healthy control and SSc samples.

Next, to examine fibroblastic state beyond numbers, gene expression changes using bulk RNA-sequencing data generated from i*Dpt;R26*^YFP^;*Csf1^fl/fl Exons^* ^5–7^ mice were analyzed. Consistent with flow cytometry results, the strongest down-regulated genes belonged to the CD206^hi^CD64+ macrophage signature (Fig. 3i, Extended data Fig. 5c). However, genes upregulated in i*Dpt;R26*^YFP^;*Csf1^fl/fl Exons^* ^5–7^ mice included fibroblast genes, notably *Dpt, Pi16*, *Pdpn*, *Pdgfra*, and *Cdh11,* and ECM-related genes such as *Col1a1* and *Col1a2* (Extended data Fig. 5c, Extended data Table 2). Interestingly, other upregulated genes included hyaluronan synthase *Has1* and *Has2* (Extended data Fig. 5d,e). As previously reported, loss of CSF1-dependent macrophages in hypodermal regions of skin can lead to an accumulation of ECM components such as hyaluronic acid (HA), as clearance is mediated via lymphatic vessel endothelial hyaluronan receptor 1 (LYVE-1)+ macrophages^5^. The CD206^hi^CD64+ population of macrophages profiled express *Lyve1* (Fig. 2i) and is downregulated in i*Dpt;R26*^YFP^;*Csf1^fl/fl Exons^* ^5–7^ mice (Extended data Fig. 5c,f, Extended data Table 2). Loss of CD206^hi^CD64+ macrophages and upregulation of HA highlights a conserved mechanism present in multiple layers of the skin.

To investigate architectural changes in the skin microenvironment due to *Csf1* and CD206^hi^CD64+ macrophage loss, along with transcriptional changes in fibroblast and ECM-related genes, full thickness samples of flank skin from control and i*Dpt;R26*^YFP^;*Csf1^fl/fl Exons^* ^5–7^ mice were collected for histopathology. Tissue structure was histologically normal with orderly organization of all skin layers and adnexal structures (Extended data Fig. 5g). However, skin thickness was modestly increased in i*Dpt;R26*^YFP^;*Csf1^fl/fl Exons^* ^5–7^ mice (Extended data Fig. 5h), predominantly due to expansion of the dermal layer (Extended data Fig. 5i). Phenotypic changes in other skin layers including epidermis, dWAT and panniculus muscle layer (Extended data Fig. 5j,k,l) were not observed.

Collectively, these results demonstrate that more complex cell circuits may be at play in maintaining skin homeostasis. Although no impact on fibroblast abundance was observed upon acute loss of *Csf1* in *Dpt*+ fibroblasts and concomitant loss of macrophages, a modest upregulation of fibroblast- and ECM-related genes and changes to skin architecture were observed.

### Fibroblast-derived *Csf1* is required for efficient wound healing

Our current findings led us to examine how loss of fibroblast-derived *Csf1*, loss of CD206^hi^CD64+ macrophages, and baseline phenotypic changes in skin would impact wound healing in an experimental model of skin wounding in control and i*Dpt;R26*^YFP^;*Csf1^fl/fl Exons^* ^5–7^ mice. Using scRNA-seq data from Hu *et al*.^7^ which profiled different cell types involved in wound healing in mouse skin and identified the emergence and increased abundance of CD206+ myeloid cells during intermediate and late-stages (after Day 3) of wound healing, gene sets were derived from these early, intermediate, and late stage time points. These wound healing signatures were scored across the previously defined CD206^hi^CD64+ and CD11c+ macrophage populations (Fig. 4a,b). Consistent with previous results, a preferential enrichment of the late wound healing gene signature^7^ in CD206^hi^CD64+ macrophages was observed. This wound healing gene signature was also reduced in skin of i*Dpt;R26*^YFP^;*Csf1^fl/fl Exons^* ^5–7^ compared to control mice profiled by bulk RNA-seq (Fig. 4c,d, Extended Data Fig. 6a). These data implicate *Csf1*-dependent CD206^hi^CD64+ macrophages in the wound healing response. We next wanted to determine if wound healing defects would be observed in i*Dpt;R26*^YFP^;*Csf1^fl/fl^* mice. Using a well-established unsplinted model of wound healing^29,30^, full thickness, circular wounds were generated on the dorsal skin of i*Dpt;R26*^YFP^;*Csf1^fl/fl Exons^* ^5–7^ and control mice on tamoxifen (Fig. 4e). Wound healing kinetics in i*Dpt;R26*^YFP^;*Csf1^fl/fl Exons^* ^5–7^ mice were examined. A significant delay in percent wound closure in i*Dpt;R26*^YFP^;*Csf1^fl/fl Exons^* ^5–7^ mice compared to control was observed (Fig. 4e,f, Extended Data Fig. 6b). This delay in wound closure became apparent on Day 4 post-injury, where the average percent closure was 43% in control mice compared to an average of 28% in i*Dpt;R26*^YFP^;*Csf1^fl/fl Exons^* ^5–7^ mice. Significant delays in percent closure of the wound continued to Day 7 (control: ∼71%, i*Dpt;R26*^YFP^;*Csf1^fl/fl Exons^* ^5–7^ ∼60%). By Day 9 post-injury, percent closure of the wound was comparable, with control ∼83% and i*Dpt;R26*^YFP^;*Csf1^fl/fl Exons^* ^5–7^ ∼75%. Pronounced soft tissue swelling at the wound margins of control mice at Day 2-4 were consistent with acute congestion, edema, and inflammation, which diminished at days 6-7 as the margins of the wound contracted with normal healing. By contrast, little gross evidence of an acute inflammatory response was present at the margins of the wound in i*Dpt;R26*^YFP^;*Csf1^fl/fl Exons^* ^5–7^ mice at days 2-4, with little evidence of contracture at later time points. Consistent with previous studies, these data demonstrate CD206^hi^CD64+ macrophages are essential during intermediate/late phases of wound healing (Day 4-7 post-injury). These results suggest that fibroblast-derived *Csf1* maintains a CD206^hi^CD64+ macrophage population that is required for effective wound healing.

### Fibroblast-derived *Csf1* is increased in human systemic sclerosis

Stromal cells are major drivers of fibrosis, which is typically characterized by an exaggerated wound healing response to injury or other insults. Fibrosis is a hallmark of several connective tissue diseases, including systemic sclerosis (scleroderma; SSc), which is an autoimmune disease with fibrotic manifestations across several major organs, including skin^15,31^. Interestingly, fibroblasts and macrophages have been previously implicated in contributing to the pathogenesis of SSc^32–35^; however, any dependency on a CSF1-CSF1R signaling axis in this etiology remains unclear. We next interrogated if the CD206^hi^ fibroblast-derived CSF1 axis influenced human SSc.

First, publicly-available scRNA-seq datasets^32,36^ profiling skin biopsies from healthy controls and SSc patients were examined. *CSF1* expression was detected across fibroblasts, pericytes and endothelial cells in SSc (Fig. 5a, Extended data Fig. 7a). Correlation with disease status revealed that pseudobulk expression of *CSF1* was significantly enriched (p=0.000852) in fibroblasts from SSc biopsies compared to healthy controls (Fig. 5b). Weaker associations by disease status were observed for pericytes (p=0.0159), and no association was identified for endothelial cells (p=0.206; Fig. 5b). Fibroblast-specific *CSF1* expression (Fig. 5c,d) was further elevated in a subtype of patients with diffuse-SSC (dSSC)^32^, defined by increased systemic severity (Fig. 5e). *CSF1* expression is restricted to three *DPT+* fibroblast clusters, including one that expressed *LGR5,* a previously defined scleroderma-associated fibroblast (ScAF) subtype morphologically and molecularly perturbed in SSc patients^32^ (Fig. 5f). Our findings suggest that *DPT+* fibroblasts specifically up-regulate *CSF1* in human SSc in accordance with disease severity.

To validate the presence of *CSF1*-expressing fibroblasts, RNA in situ hybridization (ISH) was performed in human skin tissue sections from an independent cohort of patients with SSc (Fig. 5g). Consistent with previous studies profiling fibroblast populations in human skin^14,37^, *PDGFRa* was used as a pan-fibroblast marker with co-staining for *CSF1*. In both control and SSc patient samples, colocalization of *PDGFRa* and *CSF1* transcripts was observed (Fig. 5g, Extended data Fig. 7b, arrowheads), consistent with observations made in single-cell datasets.

To investigate the implications of elevated *CSF1* on the macrophage compartment in SSc, correlations with macrophage abundance were computed in a cohort where sufficient CD45+ cells were sampled^36^. A strong positive correlation was observed between fibroblast-specific *CSF1* expression with the percentage of macrophages that expressed *MRC1* (CD206) and *CSF1R* (r=0.548, p<0.05; Fig. 5h,i; Extended Data Fig. 7c). In contrast, pericyte and endothelial *CSF1* showed no significant association with macrophage abundance (p>0.05, Extended Data Fig. 7d). These associations were validated in a separate human study^38^, where *CSF1* expression and a fibroblast signature (Fig. 5j-l), but not pericyte or endothelial cells (Extended Data Fig. 7f), were positively correlated with a CD206+ macrophage gene signature (*MRC1, FCGR1A, CSF1R, TIMD4* and *ADGRE1*) in both SSc samples or healthy controls samples independently (Fig. 5k), and when combined (Extended Data Fig. 7e). Our findings suggest that fibroblasts in human skin are the key cellular source of *CSF1* for skin macrophages in both healthy and patients with severe SSc.

## Discussion

Here, we developed novel fibroblast-restricted genetic tools to specifically target *Dpt*+ fibroblasts and elucidate how they support macrophage function in skin. Fibroblast and macrophage heterogeneity in the skin was extensively profiled using a combination of single-cell technologies, flow cytometry, and in situ hybridization (ISH). This combination of approaches demonstrated that a specific subset of CD206^hi^CD64+ macrophages is highly dependent on fibroblast-derived CSF1 in skin.

In mouse skin, fibroblasts are the main source of CSF1 and encompass several transcriptionally distinct clusters that reside across different histological layers^5,17^. We found that CD206^hi^CD64+ macrophages are dependent on fibroblast-derived CSF1 for their maintenance. These CD206^hi^CD64+ macrophages express markers of tissue residency^13,24^, including *Timd4, Lyve1*, and *Folr2*, and are distinguishable from other macrophage subsets in the skin using CD64 as a surface marker. CD206^hi^CD64+ macrophages reside in the dermis, dWAT, and adventitia, and likely receive CSF1 from *Dpt*+ fibroblasts co-inhabiting the same compartments. The concept of fibroblasts providing a supportive local microenvironment for cells of the monocyte/macrophage lineage has been supported by previous work. Stanley and colleagues^39^ reconstituted mice lacking *Csf1* (*Csf1*^op/op^) with a transgene that reinstated the membrane-bound isoform of CSF, showing rescue of macrophages in the dermis, indicating that macrophages in this niche require local CSF1 for homeostasis. More recently, Bajenoff and colleagues^40^ showed that endothelial-derived CSF1 in vascular beds at steady-state is required for the maintenance of non-classical monocytes, which patrol blood vessels. Aligned with these findings, our data suggests fibroblasts that express *Dpt* constitute the primary niche for skin-resident macrophages.

Previous computational and experimental approaches have modeled a fibroblast-macrophage two-cell circuit in which fibroblasts produce CSF1 to maintain macrophage growth and survival, and in turn, macrophages provide growth factors to support fibroblast proliferation^28^. While growth factors are required to maintain both populations, an investigation of cell number control mechanisms suggested that macrophages are more sensitive to growth factor availability, whereas fibroblasts are more sensitive to space limitations, and these two processes may be coupled to control cell numbers within tissues^12^. We hypothesized that CSF1 loss in *Dpt*+ fibroblasts and the accompanying reduction in CD206^hi^CD64+ macrophages may in turn impact skin fibroblasts through decreased *Igf* or *Pdgf* provision^5,28^ or increased ‘space’ within the skin microenvironment. However, in mice with conditional knockout of *Csf1* in *Dpt*+ fibroblasts, in which CD206^hi^CD64+ macrophages are markedly reduced, which may relate to levels of growth factor receptor (CSF1R) expression, no change in fibroblast abundance was observed by flow cytometry or ISH. These findings suggest that fibroblast and macrophage cell numbers are not coupled in vivo. However, the loss of *Csf1* in *Dpt*+ fibroblasts, and the resulting loss of CD206^hi^CD64+ macrophages, elicited a modest upregulation of fibroblast- and ECM-related genes, particularly *Has1* and *Has2*. Our results are consistent with a previous report showing an elevation of HA and degradation of the collagen network due a reduction in LYVE1+ hypodermal macrophages following the loss of fibroblast-derived CSF1^5^. These changes were validated at the protein level and were localized to the hypodermal adventitia^5^. The loss of *Csf1* in hypodermal fibroblasts did not lead to the differential expression of genes related to collagen or hyaluronic acid homeostasis^5^. Interestingly, we observed RNA induction of *Has1*/*Has2* in i*Dpt;R26*^YFP^;*Csf1^fl/fl Exons^* ^5–7^ mice, and these findings were accompanied by thickening of the skin, specifically in the dermal layer. Macrophages were recently shown to elicit contraction of an in silico cardiac model composed of fibroblasts and cardiomyocytes^41^, which further supports the concept that fibroblast-macrophage interactions can significantly shape tissue architecture.

Overall, the data suggest that *Dpt*+ fibroblasts support CD206^hi^CD64+ macrophages by providing an essential source of CSF1. In turn, CD206^hi^CD64+ macrophages may regulate the function of skin fibroblasts, as loss of this macrophage population leads to transcriptional and structural changes. One additional factor for consideration is the remaining CD11c+ macrophages that, in the absence of CD206^hi^CD64+ macrophages, may be driving these fibroblast- and ECM-related gene changes. As these findings are derived from bulk RNA-sequencing of whole skin, a detailed understanding of changes in transcriptional states at single cell resolution following *Csf1* conditional knockout in *Dpt*+ fibroblasts has yet to be explored.

Beyond functionally mapping the molecular interactions that maintain homeostasis of macrophages and fibroblasts, and architecture of the skin microenvironment, this cellular network was further explored in a mouse model of skin injury and fibrosis as well as in a human skin disease. The complexity and coordination of different cell types involved during the early (inflammation), intermediate (repair/growth) and late (resolution) phases of tissue repair have been described. Previous findings reported a CD206+CD301b+ subset of macrophages important for mid-stage wound healing^6^. Analysis of scRNA-seq data confirmed the CD206^hi^CD64+ population of macrophages express *Mgl2* (the gene for CD301b, Extended Data Fig. 2b). CD206^hi^CD64+ macrophages were also enriched for genes identified in late wound-associated macrophages^7^. Loss of these cells led to a significant delay in wound closure during the intermediate and late stages of wound healing, suggesting that CD206^hi^CD64+ macrophages are essential for tissue repair and are dependent on *Csf1* derived from *Dpt*+ fibroblasts. However, the ability for the wounds to resolve at the final stage of wound healing suggests more complex mechanisms at play. The residual CD206^hi^CD64+ macrophages may be sufficient to promote healing, or the remaining CD11c+ macrophages following *Csf1* loss in *Dpt*+ fibroblasts may switch to a more reparative role. Future studies should address specific changes in fibroblasts, macrophages, and CSF1 levels that regulate the wound healing process. Additionally, baseline dermal and accompanying matrix alterations may also be contributing factors, all which require further exploration.

Finally, we highlight the role of fibroblasts in providing CSF1 to macrophages in the context of human systemic sclerosis (scleroderma; SSc), an autoimmune disease that exhibits extensive tissue fibrosis^15,31^. Transcriptomic profiling of skin biopsies from SSc patients revealed fibroblasts express markedly higher levels of CSF1, which corresponds to higher disease severity and increased macrophage abundance. These data suggest fibroblasts engage with macrophages through provision of *CSF1* in SSc. In turn, macrophages can play a significant role in the pathogenesis of SSc by secreting a variety of cytokines, including TGF-β, IL6 and PDGF, which can drive fibroblast proliferation and activation, and promote their differentiation into myofibroblasts leading to excessive ECM deposition and fibrosis^42–44^. Additionally, macrophages can secrete proinflammatory cytokines such as TNF-ɑ and IL-6, and chemokines that recruit other immune cells, contributing to a chronic inflammatory environment and amplifying the immune response, furthering tissue damage^42–44^.

Therapeutic interventions that involve CSF1R blockade have faced significant challenges, including negative impacts on immune function and concerns regarding safety and tolerability^45,46^. Our observations reveal that increased levels of CSF1, primarily derived from fibroblasts, leads to an increase in macrophage numbers in SSc. Developing alternative strategies that disrupt the provision of CSF1 by fibroblasts to disease-promoting macrophages, may reduce fibrosis, improve tissue function, and mitigate disease progression. Targeting fibroblasts in SSc has already shown promise as an effective therapeutic strategy^47^. Future research is essential to refine these fibroblast-targeted strategies and understand the underlying mechanisms that contribute to pathology of the disease.

## Methods

### Mice and in vivo treatments

*Dpt*^IresCreERT2^ (*iDpt*) mice were designed, generated, and bred at Genentech^14^. *Rosa26*^LSL.YFP/DTR^ mice were bred at Genentech and The Centre for Phenogenomics (TCP) (Jackson Labs (JAX) stock numbers 006148 (YFP) and 007900 (DTR). *Csf1^fl/fl Exons^* ^4–6^ mice with the *Csf1* gene flanked by *loxP* sites (*Csf1* fl/fl) were donated by Sherry Werner (Department of Pathology, University of Texas Health Science Center, San Antonio, USA)^27^ by way of Dr. Clinton Robbins (Department of Immunology, University of Toronto) and crossed with *Cre*ERT2 and *Cre* strains at TCP resulting in *Csf1* knockout mice (*Cre* +/-;*Csf1* fl/fl or *Cre* +/+;*Csf1* fl/fl). *Csf1^fl/fl Exons^* ^5–7^ mice were generated as described below. For i*Dpt;R26*^YFP^;*Csf1^fl/fl Exons^* ^4–6^, and i*Dpt;R26*^YFP^;*Csf1^fl/fl Exons^* ^5–7^ mouse experiments, littermate *Cre* +/- or *Cre* +/+ ; *Csf1* fl/+ or *Csf1* fl/fl mice were used as controls. Male and female mice aged 6-12 weeks were used for all studies. For tamoxifen-induced cre expression, mice were injected with 2 mg tamoxifen (Sigma, cat# T5648) diluted in sunflower seed oil (Sigma, cat# 88921) one injection a day for 5 days, one injection on alternating days for one week, and one injection the day preceding euthanasia or via injection of 2 mg tamoxifen diluted in sunflower seed oil as above one injection a day for 5 days and fed chow containing Tamoxifen (Envigo, cat# TD130.859). All mouse studies were approved by the Genentech Institutional Animal Care and Use Committee (IACUC) or the Animal Care Committee (ACC) at TCP. No statistical methods were used to predetermine sample size. All experiments were not randomized, and the investigators were not blinded to allocation for the experiments and outcome assessment, with the exception for the analysis of the wound healing assay where the investigators were blinded on the background of the mice for manual annotation of wound boundaries.

### Generation of *Csf1^fl/fl Exons^* ^5–7^ mice

Mice carrying a *Csf1* conditional knockout Csf1<tm1.1Gne> allele (MGI:7713437) were generated at Genentech using C2 ES cells (C57BL/6NTac) using established methods. The targeting vector consisted of a 1,535 bp 5’ homology arm corresponding to GRCm38/mm10 chr3:107,750,673-107,752,207, a loxP site located 215 bp upstream of exon 5, an FRT-*Pgk1*-neo-FRT-loxP cassette located in intron 7, 163 bp upstream of exon 8, and a 1,724 bp 3’ homology arm corresponding to chr3:107,745,224-107,746,947. Correctly targeted ES cell clones were identified by PCR, digital PCR, and the junction PCR products were analyzed by Sanger sequencing for correct targeting. The Neo cassette was excised in ES cells using Adeno-FlpO and clones with normal karyotyping were microinjected into albino BL/6N embryos using established methods. Cell line karyotyping was carried out by Chih-Lin Hsieh (University of Southern California, Los Angeles, CA). Resulting chimeras were bred to C57BL/6N females to achieve germline transmission. N1 heterozygous pups were verified by long range PCR followed by sequence confirmation before establishing a breeding colony. Routine genotyping of mice was performed by PCR using tail DNA with the following primers: P1 (5’-ATGGTTCAGGACACTCTCTT-3’), P2 (5’-GCAGCAGCATCAGTCTTG-3’), and P3 (5’-GGGACCTCCTAGCTTACC-3’). Amplification with P1 and P2 yields a 205 bp product from the wildtype allele and a 239 bp product from the floxed allele. Amplification with P1 and P3 yields a 266 bp product from the knockout allele resulting from Cre exposure.

### Mouse tissue digestion and stromal cell isolation and/or identification by FACS

Tissues were isolated and fibroblasts and macrophage were isolated as previously described^14,48^. To isolate flank skin, hair was shaved, and adipose tissue was removed. Tissues were placed and minced in a 15-mL conical tube with 5 mL digestion medium (RPMI + 2% FBS) containing 100 mg/mL Dispase (Life Technologies, cat# 10104159001), 100 mg/mL collagenase P (Roche, cat# 11249002001), and 50 mg/mL of DNase 1 (Roche, cat# 10104159001). Tubes containing minced tissues were placed in a 37°C water bath and 3 mL fractions were removed and filtered (70 μM) into PBS supplemented with 2% FBS 3 times after 8-, 12-, and 15-min incubation periods. Afterwards, isolated cells were spun at 1,600 rpm for 2 min. Cells were blocked with Fc block (Biolegend; cat#101302 (1:1000) or BD Pharmingen; cat#553142 (1:100)) for 15 minutes at 4C. Afterwards, cells were labeled with the following monoclonal antibodies purchased from BD Biosciences or BioLegend (see Flow cytometry antibodies below) for 20-30 min, unless otherwise noted. Data were acquired on a BD FACSymphony^TM^ A5 SE, A3-I or A3-II and analyzed using FlowJo version 10.8.1 or 10.9.0.

### Flow cytometry antibodies

From BD Biosciences

- BUV395 anti-mouse CD45 (catalog number 564279) 1:300
- BUV737 anti-mouse CD11b (catalog number 612800) 1:400
- BUV737 anti-mouse CD31 (catalog number 741740) 1:400
- BUV737 anti-mouse CD31 (catalog number 612802) 1:200

From Biolegend

- APC anti-mouse F4/80 antibody (catalog number 157306) 1:800
- BV 711 anti-mouse F4/80 (catalog number 123147) 1:200
- BV 785 anti-mouse CD11b (catalog number 101243) 1:200
- BV711 anti-mouse CD206 (catalog number 141727) 1:800
- PE anti-mouse CD206 (catalog number 141706) 1:200
- PE Cyanine 7 anti-mouse Gr-1 (catalog number 108416) 1:400
- BV421 anti-mouse Ly-6C antibody (catalog number 128031) 1:200
- APC Cyanine 7 anti-mouse Ly-6C (catalog number 128026) 1:200
- PE Dazzle anti-mouse Ly-6C (catalog number 128044) 1:400
- PE Cyanine 7 anti-mouse Ly-6G (catalog number 127618) 1:200
- PE anti-mouse CX3CR1 (catalog number 149006) 1:400
- PerCP/Cyanine 5.5 anti-mouse/human CD207 (catalog number 144216) 1:200
- BV605 anti-mouse CD64 (catalog number 139323) 1:200
- BV650 anti-mouse CD11C (catalog number 117339) 1:100
- BV 510 anti-mouse I-A/I-E (catalog number 107636) 1:400
- APC anti-mouse Podoplanin (catalog number 127410) 1:200-800
- PE Cyanine 7 anti-mouse CD326/Ep-CAM (catalog number 118216) 1:200
- PE anti-mouse CD140a/PDGFRα (catalog number 135906) 1:200
- BV711 anti-mouse Sca1 (catalog number 108131) 1:200

### Ex vivo BMDM and *Dpt*+ fibroblast co-culture

Tibia and femur pairs were harvested from Tg(act-EGFP)Y01Osb/eGFP mice, decapped to expose the bone marrow, placed open side down in a PCR tube perforated by an 18.5G needle, and centrifuged at 12,000 RPM for 20 seconds. Bone marrow cells were resuspended in cold PBS-EDTA. Red blood cells were lysed by Ack Lysing Buffer (Gibco; cat#A10492-01) for 3 minutes on ice. After, cells were transferred to a 15 mL conical tube and Ack buffer was neutralized with 5 mL of RPMI containing 10% FBS (Gibco; cat#A31607-01), 1% Penicillin/Streptomycin (Gibco; Cat#15140122), and 1% L-Glutamine (Gibco; Cat#25030081), hereafter referred to as R10. Cells were counted and 1×10^6^ cells were plated in R10 containing 10 ng/mL recombinant murine M-CSF (Peprotech; Cat# 315-02) for 2 days. After 2 days, cells were washed twice with PBS (Gibco; Cat#14190144) and cultured in R10 containing 20 ng/mL CSF1 for 3 days. BMDM identity was confirmed by flow cytometry. *Dpt*+ fibroblasts were cultivated by injecting (i.p.) i*Dpt;R26*^YFP-CTRL^ and i*Dpt;R26*^YFP^;*Csf1^fl/f Exons^* ^4–6^ mice with 2 mg tamoxifen (Sigma, cat# T5648) diluted in sunflower seed oil (Sigma, cat# 88921) once a day for 5 days starting at -d14, then twice the following week starting at -d7 and -d5. Mice were euthanized at -d3 and fibroblasts were digested as previously described. Cells were incubated with Fc Block (1:1000; Biolegend; cat#101302) for 5 minutes on ice, washed with PBS, then incubated with CD45 microbeads (Miltenyi Biotec; Cat#130-052-301) for 15 minutes on ice before magnetic-activated cell sorting (MACS)-based cell separation to deplete leukocytes. Afterwards, YFP^+^, PDPN^+^ CD45^−^, CD31^−^, EpCAM^−^ fibroblasts were isolated by fluorescence activated cell sorting (FACS). 2×10^4^ *Dpt*+ fibroblasts/well were plated in a 96-well plate and cultured from -d3 to d0 in D10 media. At d0, 1×10^5^ BMDM were added to i*Dpt;R26*^YFP-CTRL^ and i*Dpt;R26*^YFP^;*Csf1^fl/f Exons^* ^4–6^ fibroblasts or plated in the presence/absence of M-CSF (Peprotech; Cat# 315-02). The 96-well plate was placed on the Incucyte SX5 (Sartorius), a Live-Cell Imaging and Analysis Instrument. BMDM count was measured over 7 days by imaging GFP+ BMDM using the phase and green fluorescence channels at 10X magnification in an incubator at 37°C with 5% CO_2_. The number of BMDM was determined by the green object count analysis parameter.

### in vitro co-culture and CSF1/CSF1-R antibody blockade

*Dpt*+ fibroblasts were isolated by harvesting the skin from PBS-treated i*Dpt;R26*^YFP;CTRL^ mice. The skin was enzymatically digested with DNAse, Liberase, and collagenase P to prepare single-cell suspensions, as described above. A cocktail of antibodies (CD45, CD31, EpCAM, PDPN) was added to each cell suspension in the appropriate dilutions. Stained cells were washed and analyzed using LSRII (Becton Dickinson, FACSDiva™ V6.1.3 software). Sorted CD45-EpCAM-CD31-PDPN+ YFP+ fibroblasts were expanded for 1 week on collagen-coated plates at 37 C in low CO2 (2%) in complete DMEM then frozen down using CryoStor CS10 (catalog number 07930) media to maintain low passage number. For experimental assays, YFP+ fibroblasts were thawed from frozen stock and allowed to recover for 1 week, and passaged once before co-culture. For experiments, 20,000 YFP+ fibroblasts were plated on collagen-coated 12-well plates (Costar) 1-2 days prior to co-culture.

To isolate monocytes from the bone marrow, the tibia and a femur each mouse was collected. Bone marrow was isolated via centrifugation^49^. Briefly, an 18 g needle was used to create a small hole at the bottom of a 0.5mL microcentrifuge tube. The tibia and femur for each mouse was placed in a 0.5mL microcentrifuge tube nested within a larger 1.5 mL Eppendorf tube. The nested tubes were centrifuged at 10,000 x g for 30 seconds. Complete transfer of the marrow from the 0.5 mL tube to the 1.5mL bottom tube was confirmed by visual inspection. The bones were discarded, and the visual pellet in the 1.5 mL Eppendorf tube was treated with 1 mL of RBC Lysis buffer for two minutes at room temperature before addition of 10 mL of RPMI (2% FCS) media. Each sample was placed through a 0.70 µm sterile filter on a 15 ml conical tube, centrifuged at 300 x g for 5 minutes, then washed with 10 mL of PBS. Following bone marrow isolation, monocytes were enriched by negative selection using the monocyte isolation kit (Miltenyi 130-100-629) according to the manufacturer’s instructions. As a control, isolated monocytes were cultured in macrophage media supplemented with 50 ng/ml recombinant M-CSF (R&D 416-ML-010/CF).

50,000 isolated monocytes were added in co-culture with fibroblasts either directly or in transwells (Costar 3401). For transwells, monocytes were added to the insert above plated fibroblasts. Cells were cultured for 7 days.

For CSF1 and CSF1R blockade, 100 ug/mL of antibody (Fisher Scientific, aCSF1: 50-562-653, aCSF1R: #50-139-3620) was added to cell cultures.

### CSF1 detection

To detect circulating CSF1, whole blood was collected in serum separator tubes and left at room temperature for a minimum of 30 minutes. Following clot formation, blood samples were centrifuged at 10,000 g for 10 min at 4 degrees. 60 μL of the serum was diluted with 60 uL of Assay Diluent, and CSF1 levels were determined using ELISA (Cat#ab199084).

### Single-cell RNA-sequencing

Skin tissue was dissociated into single-cell suspensions as described above. Cells from individual skin tissue samples were simultaneously stained with corresponding mouse TotalSeq-B hashtag index antibodies (B0251-B0260; Biolegend), FACS antibody cocktail (CD45, CD31, CD3, EPCAM and PDPN), and the CITE-Seq/Total-Seq A Mouse Universal Cocktail v1.0 (Biolegend #199901). CITE-seq staining was performed according to Biolegend’s staining recommendations at 1:40. Indexed samples were washed with MACS buffer and pooled together into a single tube for further labeling with live-dead enrichment viability dyes (Calcein Violet, catalog number C34858, and 7AAD, catalog number 420403). Cells were sorted for live stromal cell fractions (Calcein Violet+, 7AAD-, CD45-, EpCAM-, CD31+, PDPN+ and CD31-/PDPN-double negative) and myeloid fractions (7AAD-, EpCAM-, CD45+, CD3-). Cells were counted using a haemocytometer and resuspended in MACS buffer at the appropriate cell concentration for single-cell sequencing using the Chromium Single-Cell v2 3.1′ Chemistry Library, Gel Bead, Multiplex and Chip Kits (10x Genomics), according to the manufacturer’s protocol. A total of 10,000 cells were targeted per well.

### Sequencing and single-cell data processing

Gene-expression libraries were sequenced on the NovaSeq platform (Illumina) with paired-end sequencing and dual indexing, whilst hashtag libraries were sequenced on the MiSeq platform (Illumina). A total of 26, 8 and 98 cycles were run for Read 1, i7 index and Read 2, respectively. FASTQ files from gene-expression and hashtag libraries were processed using the Cell Ranger Single Cell v.7.1.0 software (10x Genomics). In brief, scRNA-Seq data were processed using CellRanger count function (10x Genomics) and mapped to the mouse reference genome mm10 that was customized to contain the YFP sequence plasmid.

Filtered gene-cell count matrices that were output from the Cell Ranger count pipeline were imported into Seurat v.4.3.043^50–54^ using R v.3.6.1 and log-normalized using the NormalizeData function. Hashtag antibody count matrices were imported into Seurat and individual samples were demultiplexed using the demuxEM v0.1.744 package. Only cells determined as “Singlets” by demuxEM were included for analysis. Dimensional reduction was performed using the SCTransform workflow^55^ built within the Seurat package, which was utilized for downstream Principal Component Analysis (RunPCA) and clustering based on shared nearest neighbors (FindNeighbors) and leiden graph-based clustering using the original Louvain algorithm (FindClusters). A total of 30 principal components (default) were used as input for clustering steps and UMAP reduction (RunUMAP).Broad cell types were first annotated using the SingleR package^56^ with the ImmGen reference gene signature database. For CITE-Seq data processing, raw antibody-derived tag (ADT) counts were normalized using the DSB method^57^ to background signals from empty-droplets.

For the skin fibroblasts, a resolution of 0.4 was used and clusters were annotated for Pi16+ adventitial fibroblasts clusters 1 and 2 (Pi16, Gpx3), Cthrc1+ myofibroblasts (Cthrc1, Acta2), Cxcl12+ dermal fibroblasts (Cxcl12, Plac8, Dcn) and Sparc+ Dermal fibroblasts (Sparc). For macrophages, a resolution of 0.4 was used and macrophages clusters were annotated as Spp1+ (*Spp1*, *Itgax*), Mki67+ cycling (*Mki67*, *Aurka*, *Aurkb*), CD206^hi^CD64+ (*Mrc1*, *Adgre1*/F480, *Timd4*/TIM-4, CD64 and *Csf1r*), and CD11c+ (*Itgam*, CD11b). The top 100 genes, ranked by fold-change, were used for gene ontology analysis with the Biological Processes sub-ontology database in the ClusterProfiler package^58–60^. Differential abundances between cell neighborhoods of PBS and DT treated conditions were computed using the Milo method^19^ and run using default parameters as recommended by the developers. Cell-cell communication predictions between healthy fibroblasts and macrophage clusters was performed using the CellChat package^26^. The top five ligand-receptor interactions as defined using p-values derived from CellChat.

### Tissue processing and Bulk-RNA sequencing

Mouse skin tissues were isolated from i*Dpt;R26*^YFP-CTRL^ and i*Dpt;R26*^YFP^;*Csf1^fl/fl Exons^* ^5–7^ mice and dissociated as described above. RNA was extracted from cell pellets using the RNAeasy Mini Kit (Qiagen). Total RNA was quantified with Qubit RNA HS Assay Kit (Thermo Fisher Scientific) and quality was assessed using RNA ScreenTape on TapeStation 4200 (Agilent Technologies). cDNA library was generated from 2 nanograms of total RNA using Smart-Seq V4 Ultra Low Input RNA Kit (Takara). 150 picograms of cDNA were used to make sequencing libraries by Nextera XT DNA Sample Preparation Kit (Illumina). Libraries were quantified with Qubit dsDNA HS Assay Kit (Thermo Fisher Scientific) and the average library size was determined using D1000 ScreenTape on TapeStation 4200 (Agilent Technologies). Libraries were pooled and sequenced on NovaSeq 6000 (Illumina) to generate 30 millions single-end 50-base pair reads for each sample.

### Bulk-RNA data processing

RNA-sequencing data were analyzed using HTSeqGenie in BioConductor^61^ as follows: first, reads with low nucleotide qualities (70% of bases with quality <23) or matches to rRNA and adapter sequences were removed. The remaining reads were aligned to the human reference genome (human: GRCh38.p10, mouse: GRCm38.p5) using GSNAP^62,63^ version ‘2013-10-10-v2’, allowing maximum of two mismatches per 75 base sequence (parameters: ‘-M 2 -n 10 -B 2 -i 1 -N 1 -w 200000 -E 1 --pairmax-rna=200000 --clip-overlap’). Transcript annotation was based on the Gencode genes data base (human: GENCODE 27, mouse: GENCODE M15). To quantify gene expression levels, the number of reads mapping unambiguously to the exons of each gene was calculated. 4.0.x: RNA-sequencing data were analyzed using HTSeqGenie in BioConductor^61^ as follows: first, reads with low nucleotide qualities (70% of bases with quality <23) or matches to ribosomal RNA and adapter sequences were removed. The remaining reads were aligned to the human reference genome (NCBI Build 38) using GSNAP^62,63^ version ‘2013-10-10’, allowing maximum of two mismatches per 75 base sequence (parameters: ‘-M 2 -n 10 -B 2 -i 1 -N 1 -w 200000 -E 1 --pairmax-rna=200000 --clip-overlap). Transcript annotation was based on the Ensembl genes data base (release 77). To quantify gene expression levels, the number of reads mapped to the exons of each RefSeq gene was calculated. For older results using NGS pipeline version 3.16 and prior we used GRCH37 and Ensembl genes data base (release 67).

### Downstream analysis of bulk-RNA sequencing

Differential expression analysis between *Dpt^IresCreERT2^Rosa26^LSLYFP^Csf1^fl/fl^* and control skin was performed using DESeq2^64^. For macrophage gene signature scoring, the top 50 differentially expressed genes from the scRNA-seq results were used and determined using the *FindAllMarkers* function in Seurat v.4.10. Signature scores were computed using the mean log-normalized expression of all genes in the signature and scaled using z-scoring. A two-sided student’s t-test was used to statistically test the difference in macrophage signatures between groups. Macrophage signatures and hallmark gene sets were also examined using gene-set enrichment analysis (GSEA) across the differentially expressed genes using the msigdbr R package^65^.

### Analysis of human systemic scleroderma single-cell and bulk RNA sequencing datasets

Processed human scRNA-Seq data from the Tabib et al 2021^36^ and Gur et al 2022^32^ studies were ingested and re-analyzed using Seurat v4.1.0. The average log-normalized expression of *CSF1* was computed for fibroblasts, pericytes and endothelial cells per individual patient across both cohorts. Macrophage abundance was determined as a percentage of all cell types sampled per individual. Pearson correlation was used to compute the relationship between the mean expression of *CSF1* in fibroblasts, pericytes and endothelial cells with macrophage abundance in *R* using the *cor* function. Statistical significance was determined using the *cor.test* function in *R*. An unpaired student’s t-test was used to test the difference between average CSF1 expression between healthy control and SSC groups. All cell cluster annotations and clinical metadata used were as defined by the original studies.

For bulk transcriptomics analyses of SSc, raw and transcripts per million (TPM) count matrices were ingested from the Skaug et al 2019^38^ clinical cohort (GSE130955). Differential expression statistics of *CSF1* expression between healthy and disease groups was performed using DESeq2^64^. Cell type signature scoring was computed using the mean expression of all genes from TPM values. CD206^hi^CD64+ macrophages were defined by *MRC1* (CD206), *FCGR1A* (CD64), *CSF1R*, *TIMD4* and *ADGRE1* (F4/80). For fibroblast, pericyte and endothelial cell gene signatures, the top five differentially expressed genes were computed from the Tabib et al 2021^36^ scRNA-Seq cohort using the *FindAllMarkers* function in Seurat v.4.10. Associations between *CSF1* expression and gene signatures were similarly computed using Pearson correlation in *R* using the *cor* and *cor.test* functions.

### Full-thickness wounding

Prior to wounding in i*Dpt;R26*^YFP-CTRL^ and i*Dpt;R26*^YFP^;*Csf1^fl/fl Exons^* ^5–7^ mice, the dorsal portion of the animal’s back (from the scapular area to the lumbar area) was shaved. Residual hair was removed using Nair and then rinsed off with sterile water. On the day of wounding following induction of anesthesia, the exposed skin was prepped with alcohol. Two symmetrical full-thickness excisional wounds using a 6 mm biopsy punch were generated besides the midline. Photographs of individual wounds were taken with a digital camera while animals were still under anesthesia. Afterward, wounds were covered with a 2-cm square of Tegaderm (3M) sterile transparent dressing. Wounds were monitored daily and wound dressings were changed every other day. Images of the wound were acquired using a digital camera. Acquired images were analyzed for percent closure on ImageJ/Fiji. The area of the wound was manually traced. To calculate the area of the selected region, a plugin from https://github.com/AlejandraArnedo/Wound-healing-size-tool/wiki was downloaded into ImageJ/Fiji and the Manual Wound_healing_size_Manual_tool was selected.

### Tissue processing and histology

Tissue samples were fixed in formalin, paraffin embedded, sectioned at approximately 4-5 µm and sections were stained with Hematoxylin and Eosin using standard protocols. Slides were scanned using the Nanozoomer S360 automated digital slide scanning platform (Hamamatsu Photonics) equipped with 20x (NA 0.75) Nikon lens and skin measurements were made from scanned slides.

### RNA in situ hybridization

In situ hybridization (ISH) was performed in i*Dpt;R26*^YFP-CTRL^ and i*Dpt;R26*^YFP^;*Csf1^fl/fl Exons^* ^5–7^ mice using the reagents and protocols (RNAscope^TM^ Multiplex Fluorescent Reagent Kit v2) from Advanced Cell Diagnostics. Briefly, mouse skin was fixed in 10% neutral buffered formalin for 24 hours, transferred to 70% ethanol, and then processed for paraffin embedding. Transverse sections at 5 μm thickness were cut and then allowed to dry in a 60°C oven for 1 hour. Sections were rehydrated with two washes of xylene for 5 minutes, followed by two washes in 100% ethanol for 2 minutes each. Next, the sections were incubated in hydrogen peroxide at room temperature for 10 minutes, boiled in antigen retrieval buffer for 15 minutes, quickly rinsed in DI water, washed in 100% ethanol for 3 minutes, and then dried at room temperature for 5 minutes. The sections were then digested with proteinase for 15 min at 40°C. After digestion, the slides were washed twice with ISH wash buffer for 1 minute each and then hybridized with probes for 2 hours at 40°C. Following hybridization, amplifications and secondary hybridization steps were completed for each probe (mouse: Vivid 650 - YFP, Vivid 570 - Fcgr1, human: Vivid 650 - PDGFRa, Vivid 570 - Csf1) according to Advanced Cell Diagnostics protocol. After the final amplification, slides were washed and counterstained with DAPI and then mounted with Cytoseal^TM^ 60 mounting media for Immunofluorescence imaging.

### Multiplex immunofluorescence and data acquisition

Imaging was performed using an Olympus VS200 Research Slide Scanner (Olympus Life Science, Waltham, MA). Whole slide scans were viewed using both native viewer, OlyVIA and in-house Genenetch gSlide Viewer. Image analysis was performed using Fiji/Image J.

### Quantification of RNA *ISH*

Whole skin images were manually masked. Areas of high autofluoresce (near hair follicles) were manually corrected. Within each mask, *Yfp* and *Fcgr1* ISH signal was summed and normalized to total mask area. a.u. stands for arbitrary unit, a unit for intensity.

### Quantification, Statistical Analysis and Data Presentation

Data analysis was performed as noted in individual figure legends. Statistical analysis and data presentation was performed using GraphPad Prism 10.1.0. Exact p-values were provided. For statistical testing in Prism, data was assumed to be normally distributed.

## Data & Code availability

Processed data and analysis code will be available upon publication. No new algorithms were developed for this manuscript. FASTQ sequencing files and processed count matrices for the single-cell and bulk RNA sequencing (10X Chromium) will be deposited to the Array Express database upon publication.

## Acknowledgements.

We thank our Genentech colleagues in the GEM lab for allele design and generation and the Genetic Analysis lab for genotyping. We also thank our colleagues in Animal Resources for technical assistance, including animal breeding. We thank multiple contributors from the Research Pathology Department Core Labs at Genentech, Inc, in particular Charles Jones in Histopathology, and the Digital Pathology Imaging group for their technical support. This work was supported by funding from the US National Institutes of Health Grant AG045040 (JXJ), Welch Foundation Grant AQ-1507 (JXJ), Canadian Institute of Health Research Project Grant 471606 (MBB) and Genentech.

## Author contributions

Conceptualization: ACV, SZW, AA, MBB, SJT; Methodology: ACV, SZW, AA, HB, WPL, JZ, CH, EEM, YC, YAY, JXJ; Mouse management: WO, ML, RA; Software, formal analysis and data curation: ACV, SZW, AA, HB, SU, JAVH, CD, EEM; Investigation: ACV, SZW, AA, MBB, SJT; Writing: ACV, SZW, AA, MBB, SJT; Visualization: ACV, SZW, AA; Supervision: ATK, WF, SM, MBB, SJT

## Supplemental Figures

**Supplementary Figure 1:**
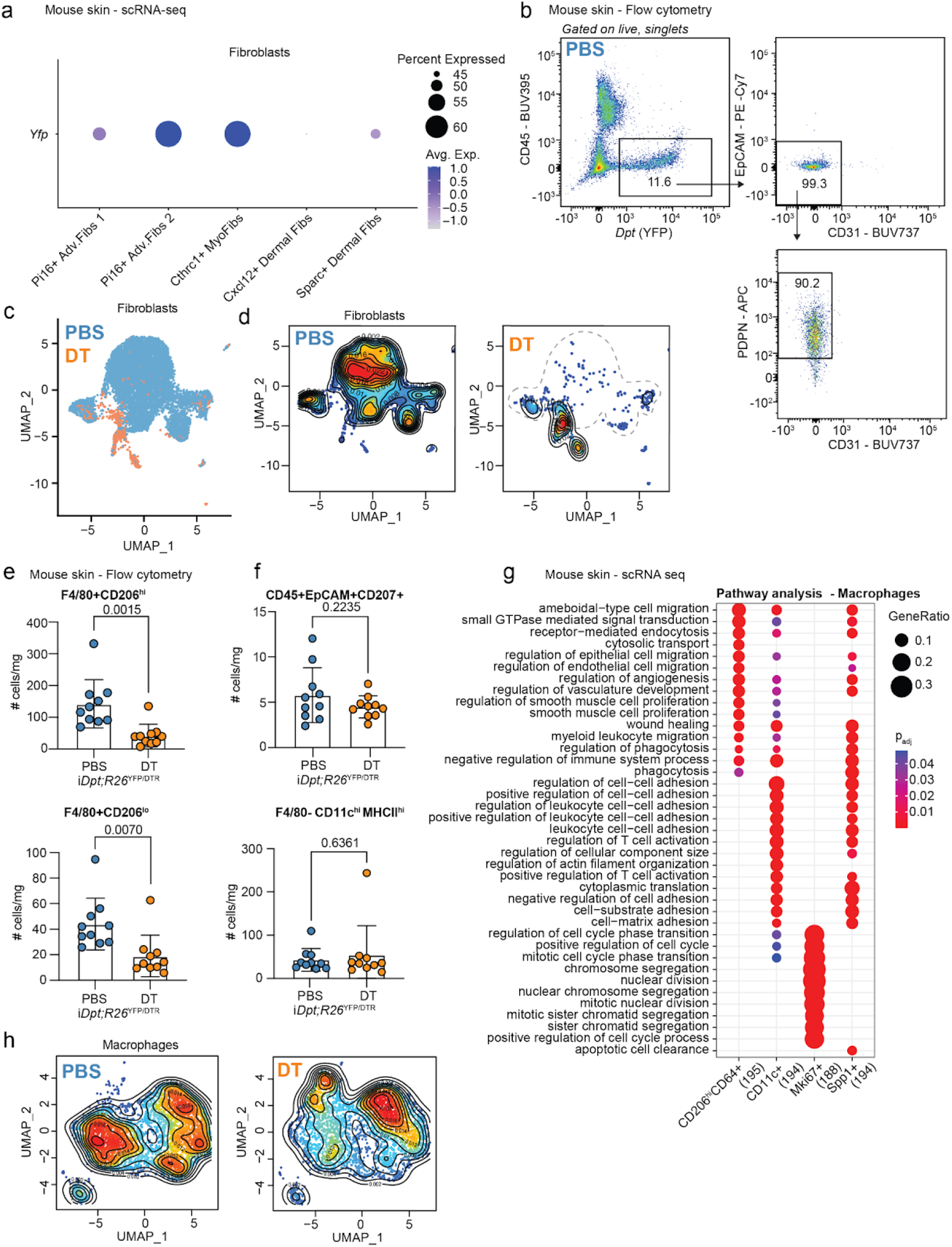
Characterization of fibroblast and macrophage compartments in i*Dpt;R26*^YFP/DTR^ mice. **a,** Single-cell sequencing in steady-state skin tissue of i*Dpt;R26*^YFP/DTR^ mice (*n*=5 PBS control, *n*=5 DT-treated, representative of 1 independent experiment). Dot plot visualizing the expression of *Yfp* across mouse fibroblasts clusters**. b,** Representative flow cytometry gating visualizing YFP expression restricted to the CD45-EpCAM-CD31-PDPN+ fibroblast compartment. **c,** UMAP visualization of fibroblast cells from scRNA-seq performed on skin from PBS and DT-treated i*Dpt;R26*^YFP/DTR^ mice. **d,** Density plots of fibroblast neighborhoods across PBS or DT conditions. (Blue, yellow, and red indicate low, medium and high density). **e,** Flow cytometry quantification of the total number of F4/80+CD206^hi^ (top) and F4/80+CD206^lo^ (bottom) cells in the skin of i*Dpt;R26*^YFP/DTR^ mice treated with PBS (*n*=10) and DT (*n*=10). Representative of 7 independent experiments. Bar plots represent Mean ± SD. Statistics were calculated using unpaired, two-tailed, Student’s *t*-test. **f,** Flow cytometry quantification of the total number of CD45+EpCAM+CD207+ (top) and CD45+Ly6G-Ly6C-F4/80-MHCII^hi^CD11C^hi^ (bottom) cells in the skin of i*Dpt;R26*^YFP/DTR^ mice treated with PBS (*n*=10) and DT (*n*=10). **g,** Enriched biological pathways between CD206^hi^CD64+, CD11c+, Mki67+, and Spp1+ macrophage clusters. **h,** Density plots of macrophage neighborhoods across PBS or DT conditions. (Blue, yellow, and red indicate low, medium and high density).

**Supplementary Figure 2:**
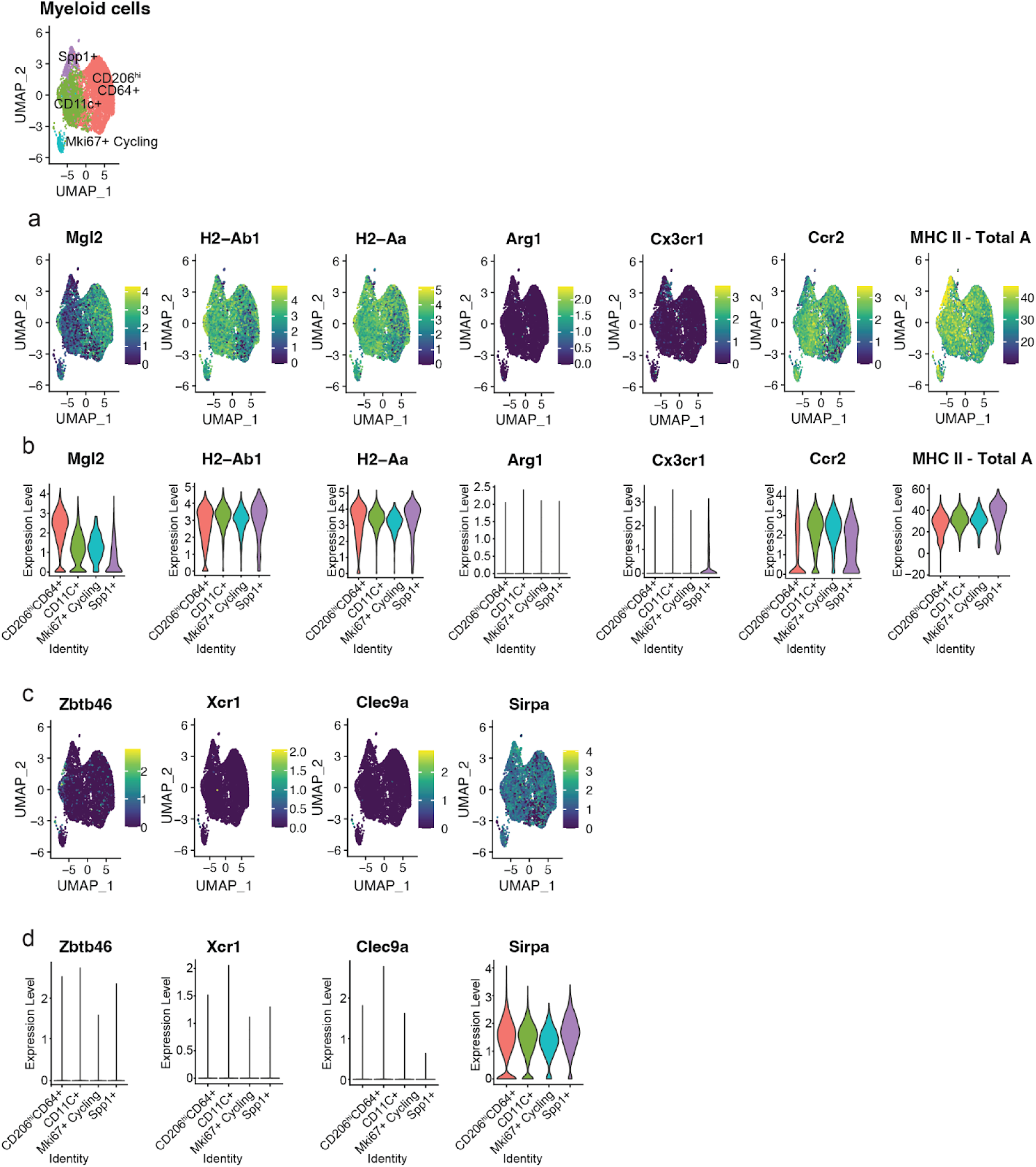
F4/80+ macrophage subsets cannot be distinguished by *MHC Class II*, *Arg1*, or *Cx3cr1* expression, and do not express markers for DCs. **a,** Feature plot of expression values for *Mgl2*, *H2-Ab1*, *H2-Aa*, *Arg1, Cx3cr1, Ccr2,* MHC Class II -Total A markers in the skin of i*Dpt;R26*^YFP/DTR^ mice. **b,**Violin plot showing expression of *Mgl2*, *H2-Ab1*, *H2-Aa*, *Arg1, Cx3cr1, Ccr2* and MHC Class II -Total A across macrophage subsets. **c,** Feature plot of expression values for *Zbtb46*, *Xcr*, *Clec9a*, *Sirpa* markers. **d,** Violin plot showing expression of *Zbtb46*, *Xcr*, *Clec9a* and *Sirpa* across macrophage subsets.

**Supplementary Figure 3:**
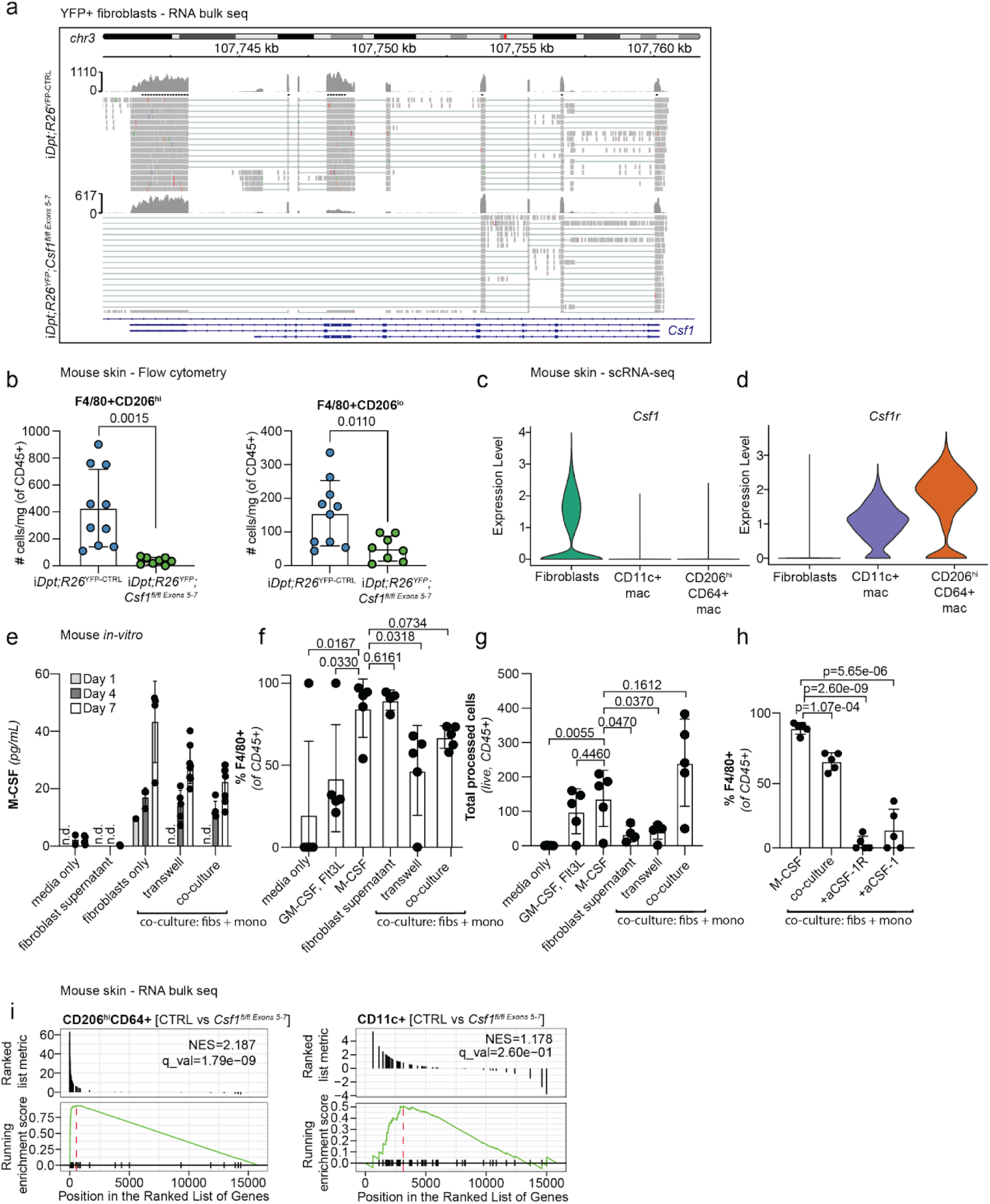
*Csf1* knockout validation in i*Dpt;R26*^YFP^;*Csf1^fl/fl Exons^* ^5–7^ mice and effect on macrophages. **a,** RNA-seq analysis of sorted YFP+ cells from the skin of control and i*Dpt;R26*^YFP^;*Csf1^fl/fl Exons^* ^5–7^ mice. Integrative Genomics Viewer (IGV) browser track displays raw sequencing reads at the *Csf1* locus. n=4 mice per group were pooled. Representative of 1 independent experiment. **b,** Flow cytometry data showing quantification of total number of CD11b+F4/80+CD206^high^ (left) and CD11b+F4/80+CD206^lo^ (right) cells from i*Dpt;R26*^YFP-CTL^ (n=10) and i*Dpt;R26*^YFP^;*Csf1^fl/fl Exons^* ^5–7^ (n=8) mice. Bar plots represent Mean ± SD. Statistics were calculated using unpaired, two-tailed, Student’s *t*-test. **c,d.** Expression of **(c)** *Csf1* and **(d)** *Csf1r from* scRNA-seq analysis across fibroblasts and CD11c+ or CD206^hi^CD64+ macrophages in the skin of i*Dpt;R26*^YFP/DTR^ mice. **e,** YFP+ fibroblasts isolated from the skin of i*Dpt;R26*^YFP-CTL^ mice co-cultured with BM monocytes (*n*=5 mice per condition, representative of 2 independent experiments). Culture supernatant was sampled one day after culture (Day 1) and on Day 4 and Day 7 for levels of M-CSF. The concentrations (pg/mL) were analyzed by Luminex. Data representative of 2 independent experiments with 3-5 replicates each. Bar plots represent Mean ± SD. Statistics were calculated using unpaired, two-tailed, Student’s *t*-test. **f,** Quantification of flow cytometry data showing the percentage of F4/80+ cells each condition on Day 7. Cells were gated on CD45+Ly6G-. Bar plots represent Mean ± SD. Statistics were calculated using unpaired, two-tailed, Student’s *t*-test. **g,** Quantification of flow cytometry data showing Total CD45+ cells in each condition. **h,** Quantification of flow cytometry data showing the percentage of F4/80+ cells in each condition with addition of CSF-1R and CSF-1 blocking antibodies. **i,** Gene-set enrichment analysis (GSEA) plot showing enrichment of the CD206^hi^CD64+ (left) and CD11c+ (right) macrophage single-cell gene signature (top 50 genes) in differentially expressed genes in i*Dpt;R26*^YFP-CTRL^ (n=7) compared to i*Dpt;R26*^YFP^;*Csf1^fl/fl Exons^* ^5–7^(n=6) skin. Individual genes of the signature are indicated by individual lines (top). Differential expression of genes (x-axis) were analyzed using DESeq2. Normalized enrichment scores (NES) and false-discovery rates (q_val) were computed using the clusterProfiler package. The leading edge for the running enrichment score is indicated by the red-line.

**Supplementary Figure 4:**
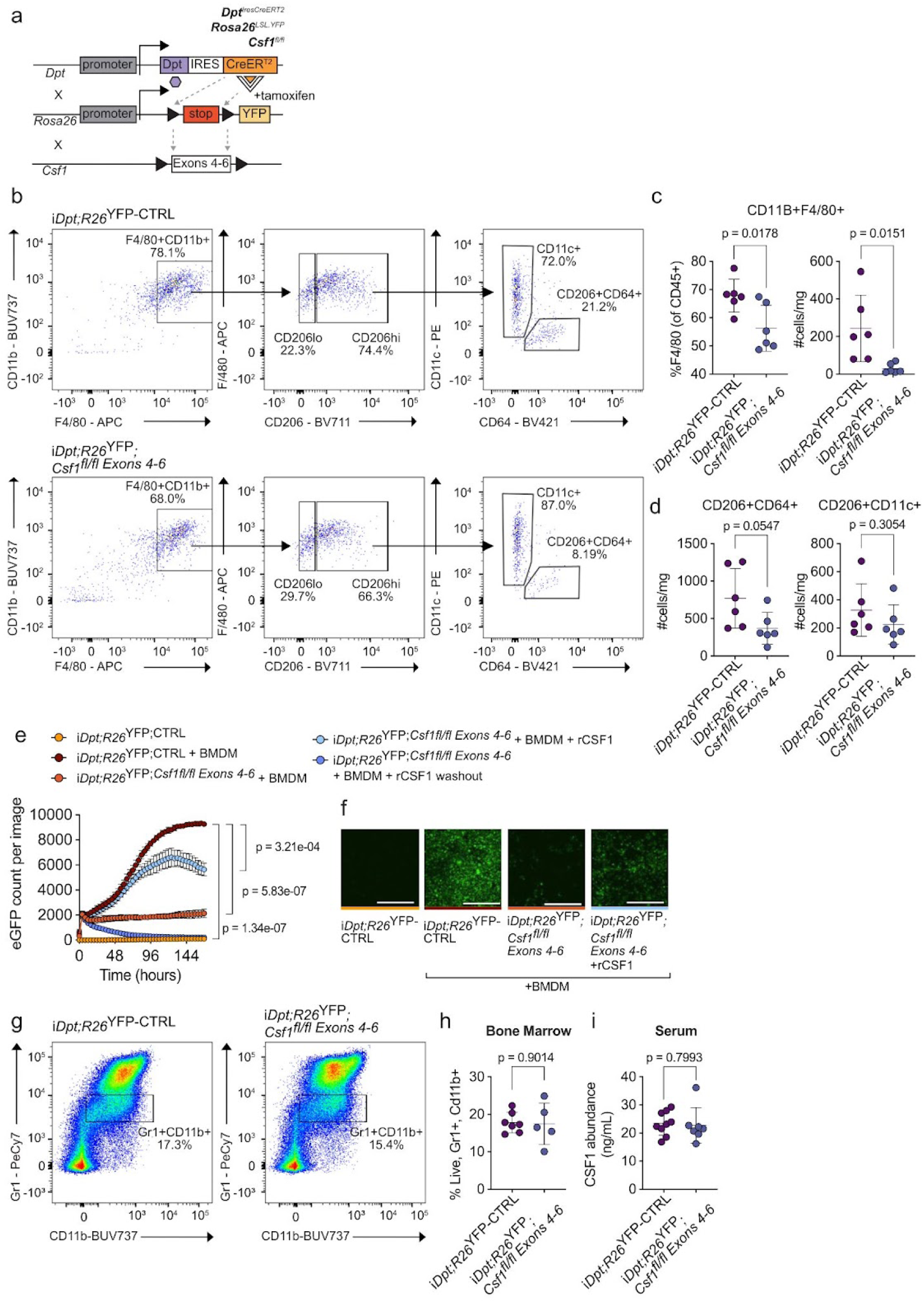
Cross-institutional validation of phenotypes observed in i*Dpt;R26*^YFP^;*Csf1^fl/fl Exons^* ^4–6^ mice. **a,** Schematic of i*Dpt;R26*^YFP^;*Csf1^fl/fl Exons^* ^4–6^ transgenic mouse. **b,** Representative gating of F4/80+CD11b+ macrophages, CD206 high and low macrophages, and CD206+CD64+ or CD206+CD11c+ macrophages in the skin of i*Dpt;R26*^YFP-CTRL^ and i*Dpt;R26*^YFP^;*Csf1^fl/fl Exons^* ^4–6^ mice after 2-weeks of tamoxifen (i.p.) administration. **c,** Quantification of percent CD11b+F4/80+ cells (left) and total number of CD11b+F4/80+ cells per mg of tissue (right) in the skin of i*Dpt;R26*^YFP-CTRL^ (n = 6) and i*Dpt;R26*^YFP^;*Csf1^fl/fl Exons^* ^4–6^ (n = 6). **d,** Quantification of total number of F4/80+CD206+CD64+ (left) and F4/80+CD206+CD11c+ cells in the skin of i*Dpt;R26*^YFP-CTRL^ (n = 3) and i*Dpt;R26*^YFP^;*Csf1^fl/fl Exons^* ^4–6^ (n = 3). **e,** Quantification of eGFP count over time in five technical replicates per condition. **f,** Representative images of eGFP BMDM ex vivo **g,** Representative gating of bone marrow Gr1+/CD11b+monocytes. **h,** Quantification of the frequency of bone marrow Gr1+/CD11b+ monocytes in i*Dpt;R26*^YFP-CTRL^ and i*Dpt;R26*^YFP^;*Csf1^fl/fl Exons^* ^4–6^ mice after 2-weeks of tamoxifen (i.p.) administration. **i,** Quantification of serum CSF1 abundance in i*Dpt;R26*^YFP-CTRL^ and i*Dpt;R26*^YFP^;*Csf1^fl/fl Exons^* ^4–6^ mice after 2-weeks of tamoxifen (*i.p.*) administration. **b,c,d** Cells were gated on Live, CD45+Gr1-cells. **g** Cells were gated on Live. Statistics were calculated using an unpaired, two-tailed, Student’s *t*-test. Data are mean ± s.d. and pooled from 2 independent experiments (**c, d, g, h, i**) or representative of 2 (**e, f**) independent experiments.

**Supplementary Figure 5:**
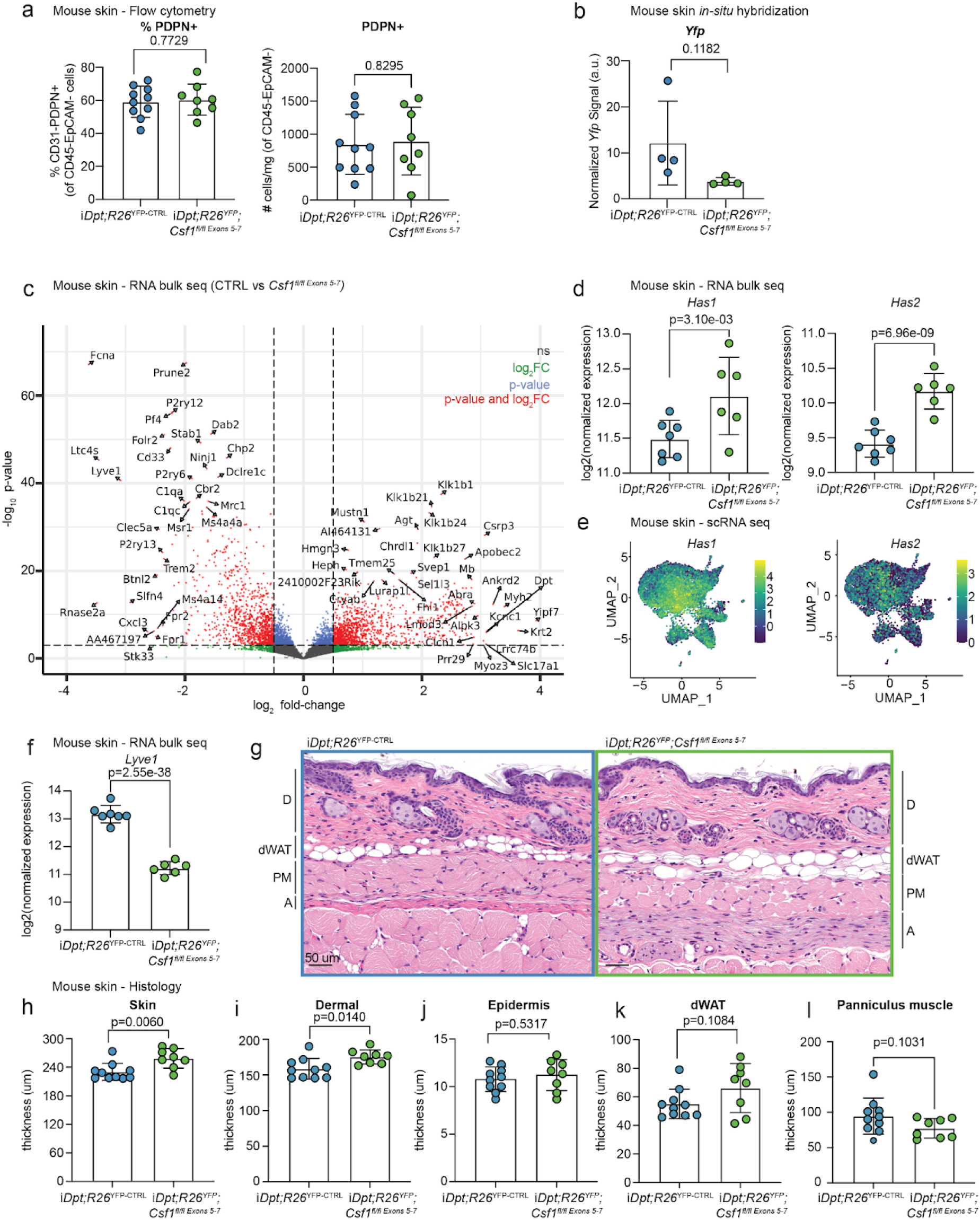
Transcriptional and phenotypic changes in the skin of i*Dpt;R26*^YFP^;*Csf1^fl/fl Exons^* ^5–7^ mice. **a,** Flow cytometry data showing quantification of the frequency (left) and total number of PDPN+CD31-cells (right) from i*Dpt;R26*^YFP-CTL^ (n=10) and i*Dpt;R26*^YFP^;*Csf1^fl/fl Exons^* ^5–7^ (n=8) mice. Bar plots represent Mean ± SD. Statistics were calculated using unpaired, two-tailed, Student’s *t*-test. **b,** in situ hybridization (ISH) quantification for *Yfp* (*Dpt*) from the skin of control (*n*=4) and i*Dpt;R26*^YFP^;*Csf1^fl/fl Exons^* ^5–7^ (*n*=4) mice. Representative of 1 independent experiment). **c,** Analysis of bulk-RNA sequencing from the skin of i*Dpt;R26*^YFP-CTL^ (*n*=7) and i*Dpt;R26*^YFP^;*Csf1^fl/fl Exons^* ^5–7^ (*n*=6) mice. Volcano plot of differentially expressed genes in the skin between i*Dpt;R26*^YFP-CTL^ and i*Dpt;R26*^YFP^;*Csf1^fl/fl Exons^* ^5–7^ mice. DESeq2 genes significantly enriched in control and conditional knockout groups are indicated by negative (left) and positive (right) fold change, respectively. **d,** Normalized expression (Log2) of hyaluronic acid related genes (*Has1* and *Has2*) in skin tissue of i*Dpt;R26*^YFP-CTL^ and i*Dpt;R26*^YFP^;*Csf1^fl/fl Exons^* ^5–7^ mice. Bar plots represent Mean ± SD. Statistical significance as determined using DESeq2. **e,** Analysis of single-cell sequencing in steady-state skin tissue of i*Dpt;R26*^YFP/DTR^ mice (*n*=5 PBS control, *n*=5 DT-treated, representative of 1 independent experiment). Feature plot of expression values for *Has1* and *Has2* in fibroblasts. **f,** Normalized expression (Log2) of hyaluronic acid receptor *Lyve-1* in skin tissue of i*Dpt;R26*^YFP-CTL^ and i*Dpt;R26*^YFP^;*Csf1^fl/fl Exons^* ^5–7^ mice. Bar plots represent Mean ± SD. Statistical significance as determined using DESeq2. **g,** Representative image of full thickness flank skin sections examined in control (n=10) and i*Dpt;R26*^YFP^;*Csf1^fl/fl Exons^* ^5–7^ mice (n=8), comprising dermis (D), dWAT (d), panniculus muscle (PM), and adventitia (A). Scale bars, 50 um. **h,i,j,k,l,** Skin thickness measurements (um) made on scanned slides in gSlideViewer. Bar plots represent Mean ± SD. Statistics were calculated using unpaired, two-tailed, Student’s *t*-test. **(h)** Overall skin thickness (um) measurements extend from the epidermal surface to the deep margin of the dWAT layer and include **(i)** dermis, **(j)** epidermis, and **(k)** dWAT. **(l)** Measurement of panniculus muscle thickness (um).

**Supplementary Figure 6:**
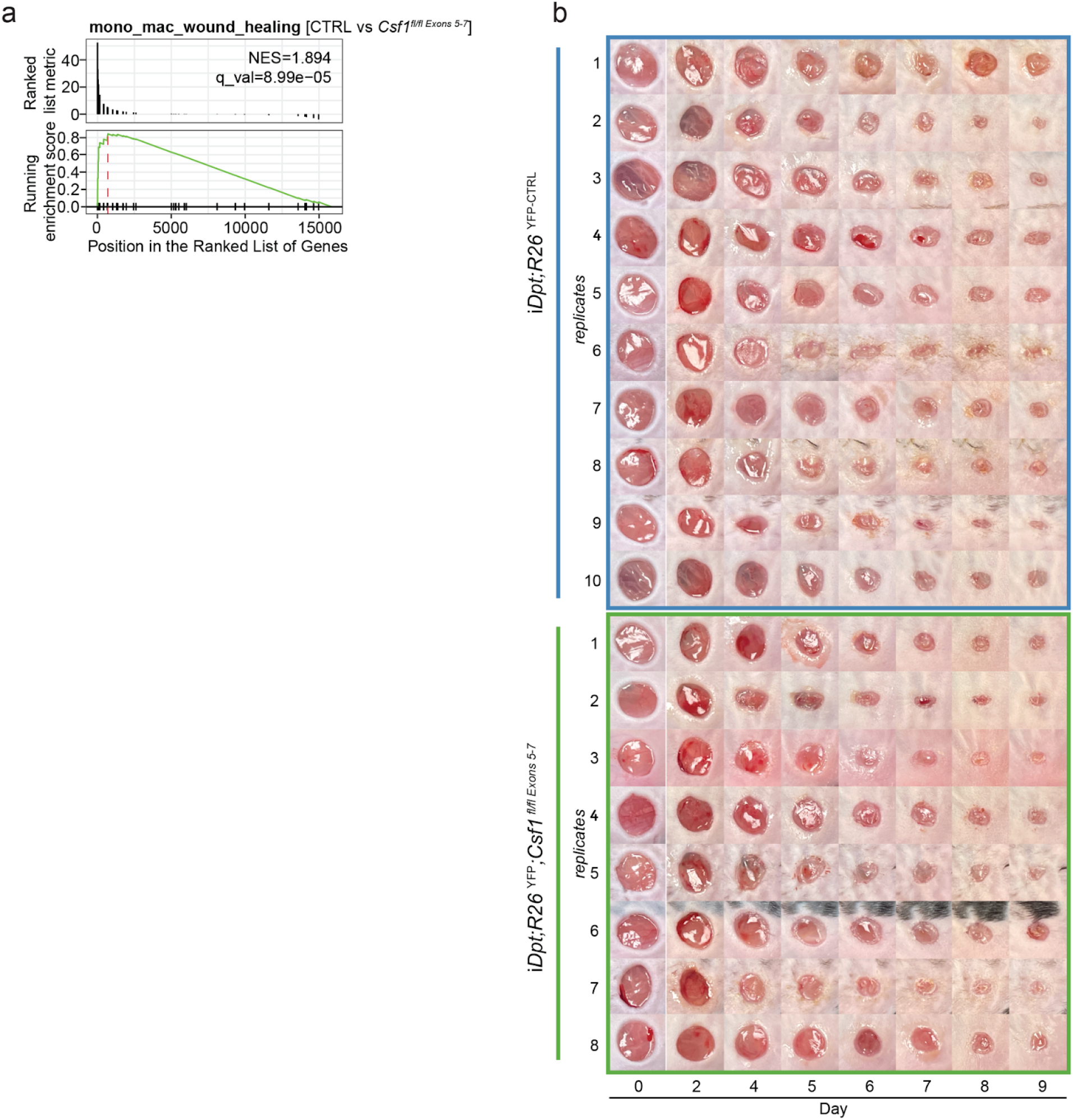
Loss of fibroblast-derived *Csf1* alters skin wound healing. **a,** Gene-set enrichment analysis (GSEA) plot showing enrichment of a macrophage wound healing gene signature from Hu et al 2023 in differentially expressed genes in i*Dpt;R26*^YFP-CTRL^ (n=7) compared to i*Dpt;R26*^YFP^;*Csf1^fl/fl Exons^* ^5–7^ (n=6) skin. Individual genes of the signature are indicated by individual lines (top). Differential expression of genes (x-axis) were analyzed using DESeq2. Normalized enrichment scores (NES) and false-discovery rates (q_val) were computed using the clusterProfiler package. The leading edge for the running enrichment score is indicated by the red line. **b,** Images of wounds from one representative experiment of both i*Dpt;R26*^YFP-CTRL^ (n=10) and i*Dpt;R26*^YFP^;*Csf1^fl/fl Exons^* ^5–7^ (n=8) mice were acquired using a digital camera. Acquired images were analyzed for percent closure on ImageJ.

**Supplementary Figure 7:**
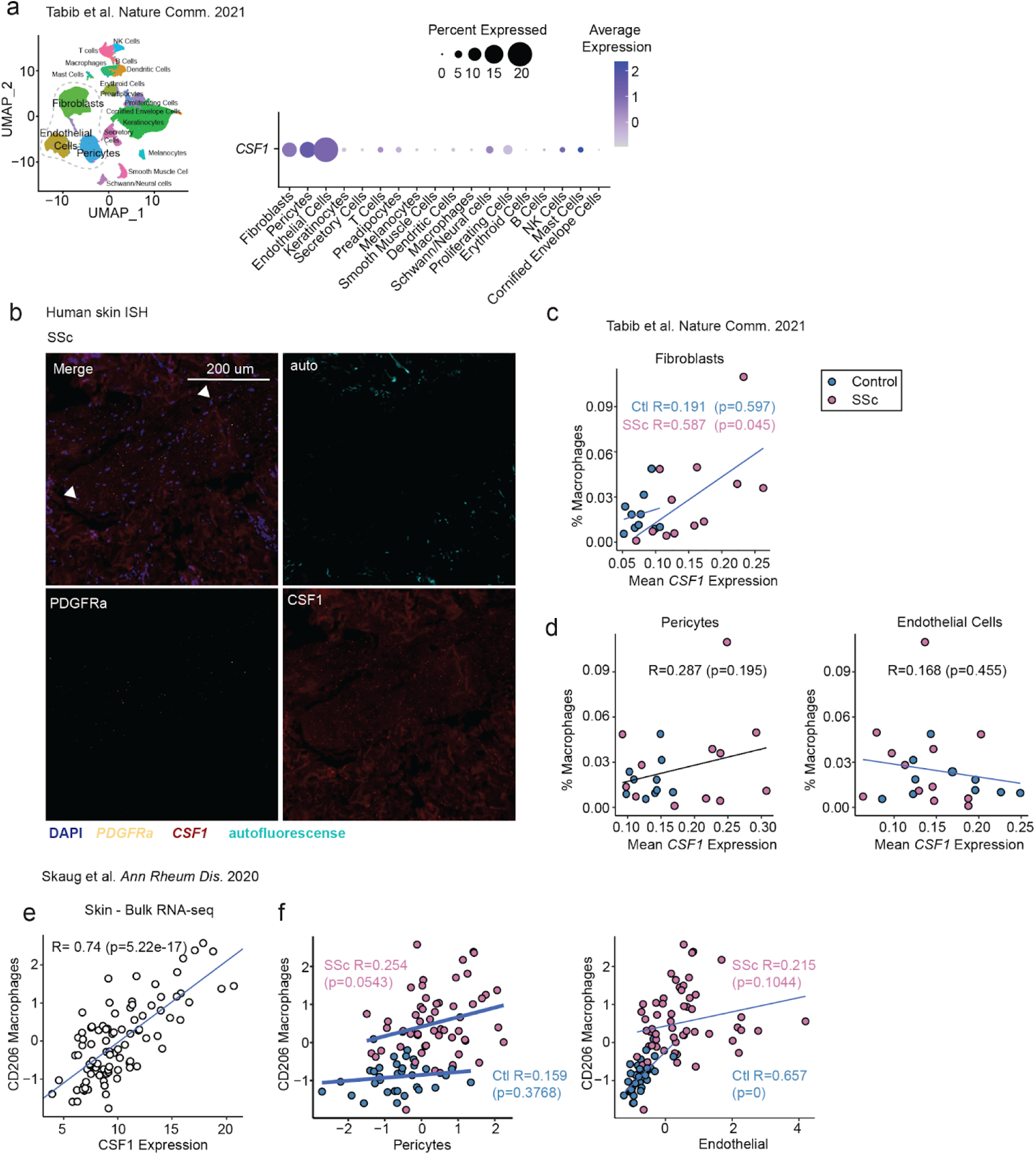
Human skin fibroblast *CSF1* correlates with macrophages. **a,** Data re-analyzed from the Tabib 2021 study^36^. DotPlot (right) showing the expression of *CSF1* across different cell clusters (left) of human scleroderma skin tissue. DotPlot indicates the average expression and abundance of *CSF1* expression across clusters, including enrichment in stromal cell populations. **b,** Representative in situ hybridization (ISH) images for *PDGFRa* (yellow) and *CSF1* (red) from human SSc skin (n=3) tissue sections from an independent cohort of patients. Representative of 1 independent experiment. DAPI signal is shown in blue. Autofluorescence is shown in cyan. Scale bars, 200 um. **c,** Correlation plot of the average *CSF1* expression in fibroblasts and the percentage of macrophages of all cells per patient group (control, R=0.1909, SSc, R=0.5872) re-analyzed from the Tabib 2021 study^36^. **d,** Correlation plot of the average *CSF1* expression in pericytes (left) and endothelial cells (right) and the percentage of macrophages of all cells per individual patient. Computed using Pearson correlation. For each Pearson correlation, statistical significance was determined using the *cor.test* function in *R.* Healthy (blue; *n*=10) and scleroderma, SSc (pink; *n*=12) samples are colored. **e,** Pearson correlation between the normalized expression of *CSF1* (TPM) and CD206 macrophage gene signature scores across all healthy control and SSc samples from the Skaug 2020 study^38^. **f,** Pearson correlation between CD206 macrophages with pericyte (left) and endothelial (right) gene signature scores in both healthy control and SSc samples.

